# Smooth muscle-derived adventitial progenitor cells promote key cell type transitions controlling plaque stability in atherosclerosis in a Klf4-dependent manner

**DOI:** 10.1101/2023.07.18.549539

**Authors:** Allison M Dubner, Sizhao Lu, Austin J Jolly, Keith A Strand, Marie F Mutryn, Tyler Hinthorn, Tysen Noble, Raphael A Nemenoff, Karen S Moulton, Mark W Majesky, Mary CM Weiser-Evans

## Abstract

We previously established that vascular smooth muscle-derived adventitial progenitor cells (AdvSca1-SM) preferentially differentiate into myofibroblasts and contribute to fibrosis in response to acute vascular injury. However, the role of these progenitor cells in chronic atherosclerosis has not been defined. Using an AdvSca1-SM lineage tracing model, scRNA-Seq, flow cytometry, and histological approaches, we confirmed that AdvSca1-SM cells localize throughout the vessel wall and atherosclerotic plaques, where they primarily differentiate into fibroblasts, SMCs, or remain in a stem-like state. Klf4 knockout specifically in AdvSca1-SM cells induced transition to a more collagen-enriched myofibroblast phenotype compared to WT mice. Additionally, Klf4 depletion drastically modified the phenotypes of non-AdvSca1-SM-derived cells, resulting in more contractile SMCs and atheroprotective macrophages. Functionally, overall plaque burden was not altered with Klf4 depletion, but multiple indices of plaque vulnerability were reduced. Collectively, these data support that modulating the AdvSca1-SM population confers increased protection from the development of unstable atherosclerotic plaques.

## INTRODUCTION

Atherosclerosis is a complex inflammatory condition and the major driver of cardiovascular disease, a spectrum of diseases resulting in approximately 32% of global deaths^1, 2^. Management of atherosclerosis involves lipid-lowering and anti-inflammatory medications. Unfortunately, statins fail to fully resolve CVD risk and, despite intense lipid lowering, many patients have residual risks. Additionally, anti-inflammatory treatments have limitations for cost and need for chronic use, which carry increased risk of infection and sepsis. Thus, additional therapeutic approaches are needed to directly target the vascular wall cells in atherosclerotic lesions^3–8^. Surprisingly, few if any therapies directly focus on the pathological mechanisms of resident vascular cells, which may provide insights for future therapy beyond current care.

Historically, research in the field of atherosclerosis has focused on the role of the innermost layer of the blood vessel, the intima, in driving atherosclerosis progression through endothelial dysfunction, lipid accumulation/oxidation, and macrophage infiltration^1, 3–5^. More recently, lineage tracing studies in murine models and human atherosclerotic tissues have defined the important role for vascular smooth muscle (SMC) in disease progression. However, while less frequently studied, it has been well recognized that cells in the outermost layer of the vessel, the adventitia, are critical contributors to early stages in the pathogenesis of atherosclerosis. Further, the “outside-in” hypothesis posits that expansion of adventitial microvessels, the vasa vasorum (VV), acts as an early and potent driver of plaque progression by facilitating inflammatory cell infiltration, tertiary lymphoid organ, and supplying the oxygen/nutrient needs of the growing plaque^6–10^.

Recent research has demonstrated that the adventitia is highly dynamic and home to a wide variety of cells, including fibroblasts, leukocytes, and resident vascular progenitor cells^7, 11–13^. Our group discovered a subpopulation of these adventitial progenitor cells (AdvSca1-SM cells) derived from mature, contractile smooth muscle cells (SMCs) that undergo KLF4-dependent reprogramming^14^. Subsequent research demonstrated that maintenance of the AdvSca1-SM stemlike phenotype is dependent on continuous KLF4 activity and that AdvSca1-SM cells preferentially differentiate into myofibroblasts to contribute to perivascular fibrosis in response to acute vascular injury^15, 16^. However, the contribution of AdvSca1-SM cells to chronic diseases such as atherosclerosis has not been established.

In this study, we characterized the multifaceted contributions of AdvSca1-SM cells to atherosclerosis. Immunofluorescence microscopy demonstrated that AdvSca1-SM-derived cells are found throughout the adventitia, media, and atherosclerotic plaque. Using single cell RNA sequencing (scRNA-Seq), we established the major differentiation trajectories of AdvSca1-SM cells, as well as their ability to communicate with other cells in the plaque microenvironment. Surprisingly, we found that AdvSca1-SM cell specific knockout of KLF4 altered both the differentiation trajectories of AdvSca1-SM cells as well as the transcriptomic profiles of SMCs and macrophages, leading to a more stable plaque phenotype. These findings indicate that AdvSca1-SM cells are major regulators of atherosclerotic plaque progression and that KLF4-dependent phenotypic transitions of both AdvSca1-SM cells and other vascular and plaque cells are critical to plaque complexity, thus highlighting a potential future therapeutic target.

## RESULTS

### Induction of atherosclerosis in AdvSca1-SM cell lineage mice

Our previous research indicated a preferential differentiation of AdvSca1-SM cells into myofibroblasts in the setting of unilateral carotid ligation, a well-established model for acute vascular injury, neointima formation, and adventitial remodeling^15, 16^. In these experiments, AdvSca1-SM cells were integral to adventitial remodeling and arterial fibrosis. However, the role of AdvSca1-SM cells in chronic vascular diseases, specifically atherosclerosis, has not been established. To elucidate the role of these adventitial progenitor cells in atherosclerosis, we utilized the AdvSca1-SM cell lineage tracing mouse model we previously developed (Gli1-Cre^ERT^/Rosa26-YFP)^15^. Upon tamoxifen injections, AdvSca1-SM cells are selectively and permanently labeled with the YFP reporter and there is a complete lack of medial SMC labeling. Following tamoxifen, AdvSca1-SM lineage mice were injected with a gain-of-function mutant PCSK9-AAV to knock down LDL receptors and placed on a high fat/high cholesterol diet for 8 to 28 weeks to induce hypercholesterolemia and atherosclerotic plaque formation (Figure 1A, Extended Data Figure 2)^17^.

**Figure 1.**
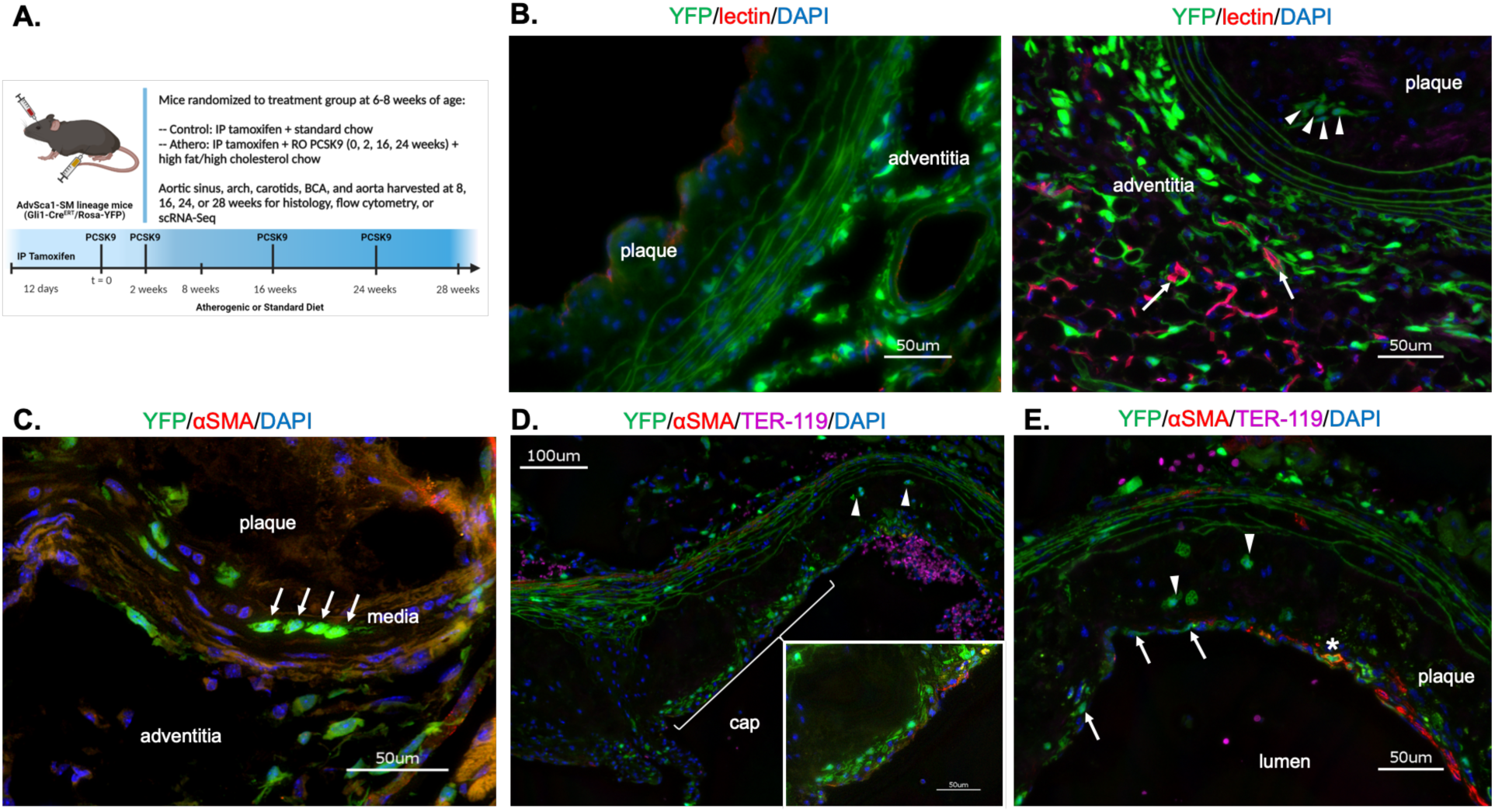
AdvSca1-SM cells are distributed throughout the vascular wall and atherosclerotic plaque. **A.** Schematic of experimental approach. **B.** A subset of animals were injected intravenously with fluorescently labeled *Griffonia simplicifolia* lectin I (GSL I) isolectin B4 5 minutes prior to sacrifice to label functional vasculature *in vivo*. Representative immunofluorescence of aortic root sections from 24-week plaques stained for YFP (AdvSca1-SM and AdvSca1-SM-derived cells; green); lectin (red); DAPI for all cell nuclei (blue). Arrows indicate functional adventitial microvasculature (lectin labeled) surrounded by YFP^+^ cells; arrowheads indicate intraplaque YFP^+^ cells. **C.** Representative immunofluorescence image of 28-week aortic root plaque stained for αSMA (red) and YFP (green). Arrows indicate YFP^+^ medial cells. **D.** and **E.** Aortic root slides from 24-week plaques stained for YFP (green), αSMA (red), Ter-119 (magenta), and DAPI (blue). Low power and high-power insert (D) show YFP^+^ cells in the cap of the plaque. Panel E shows YFP^+^ cells in the plaque cap (arrows), YFP^+^/αSMA^+^ cells in the plaque cap (asterisk), and YFP^+^ cells in the body of the plaque (arrowheads). Scale bars for all immunofluorescence images are 50 μm, except low power panel D (100 μm).

### AdvSca1-SM-derived cells are found throughout the vessel wall and plaque in both early and late stage atherosclerosis

We first sought to determine the spatial distribution of AdvSca1-SM cells and their progeny within the vessel wall and plaque. The majority of published studies of atherosclerosis progression primarily focused on lipid accumulation and monocyte recruitment on the intimal surface of the vessel^1, 3–5^. However, additional work has demonstrated the integral role of the vasa vasorum in driving plaque progression^1, 3–9^. Moreover, using an *in vivo* Matrigel^TM^ plug assay, we previously demonstrated the ability of AdvSca1-SM cells to form microvasculature, raising the question of whether AdvSca1-SM cells could be involved in the expansion of the adventitial microvasculature in the setting of atherosclerosis^14^. To investigate this possibility, a subset of mice after 16 or 24 weeks of atherogenic diet were intravenously injected with fluorescently conjugated isolectin B4 to label functional vasculature. Using this approach, we detected microvessels in the adventitial area of the aortic root. Similar to our findings in acute vascular injury, there was an expansion of YFP^+^ AdvSca1-SM cell-derived cells in the adventitia, with many frequently associated with the adventitial microvasculature, suggesting a neovascularization role for AdvSca1-SM cells in atherosclerosis and supporting our previous Matrigel^TM^ plug experiments (Figure 1B). While the majority of YFP^+^ cells were found in the adventitia, they were also observed in the medial layer of the aortic root (Figure 1C). Finally, we observed YFP^+^ cells localized in the plaque itself, both via formation of the fibrous cap (Figure 1D & 1E), as well as in the core of the plaque. These data indicate significant roles for AdvSca1-SM cells throughout the vascular wall in the context of chronic atherosclerosis and suggest multifaceted contributions to atherogenesis.

### Single cell RNA sequencing defines shifts in vascular cell phenotypes in atherosclerosis

Given the wide distribution of AdvSca1-SM cell-derived YFP^+^ cells in the vascular adventitia, media, and atherosclerotic plaque, we used unbiased single cell RNA sequencing (scRNA-Seq) to define the possible functions of AdvSca1-SM cells in atherosclerosis. As described in the Methods, arteries from AdvSca1-SM lineage tracing mice at both baseline (after tamoxifen treatment) and after 16 weeks of either control or atherogenic conditions were harvested, processed into single cell suspensions, FACS sorted based on YFP expression, and YFP^+^ and YFP^-^ cells subjected to scRNA-Seq. Datasets from all samples were combined, passed through quality control measures, and processed using the Seurat pipeline, generating a total of 26 cell clusters (Figure 2A, Extended Data Figure 4). Gene expression profiles were used to assign identities to the different clusters, and the top 5 differentially expressed genes per cluster are shown in Figure 2B. As expected, we detected shifts in cell populations between baseline and 16 weeks of control or atherogenic treatment (Figure 2C). We then examined the changes in cell populations between mice on control or atherogenic diet after 16 weeks of treatment. The clusters that showed the most variability as a result of atherosclerosis were the fibroblast clusters (Fib_1, Fib_2, Fib_3, Fib_4). Specifically, we observed increases in the size of Fib_1, Fib_3, and Fib_4 clusters, along with a decrease in Fib_2 (Figure 2D). We also observed a slight decrease in the AdvSca1-SM cell cluster, suggesting an increased pattern of differentiation into other cell types in the setting of atherosclerosis. As AdvSca1-SM cells are multipotent progenitor cells and don’t represent a static cell population, RNA velocity analysis was used to predict future cell transition patterns, as described previously^18^. Validating our previous findings, the streamplots indicate significant differentiation away from the most stemlike state, the AdvSca1-SM cell cluster, and into other cell types (Figure 2E). These data are in agreement with previous scRNA-Seq data showing shifts in cell phenotype with atherosclerosis progression and confirm a transition of AdvSca1-SM cell phenotype away from a stemlike phenotype^19–23^.

**Figure 2.**
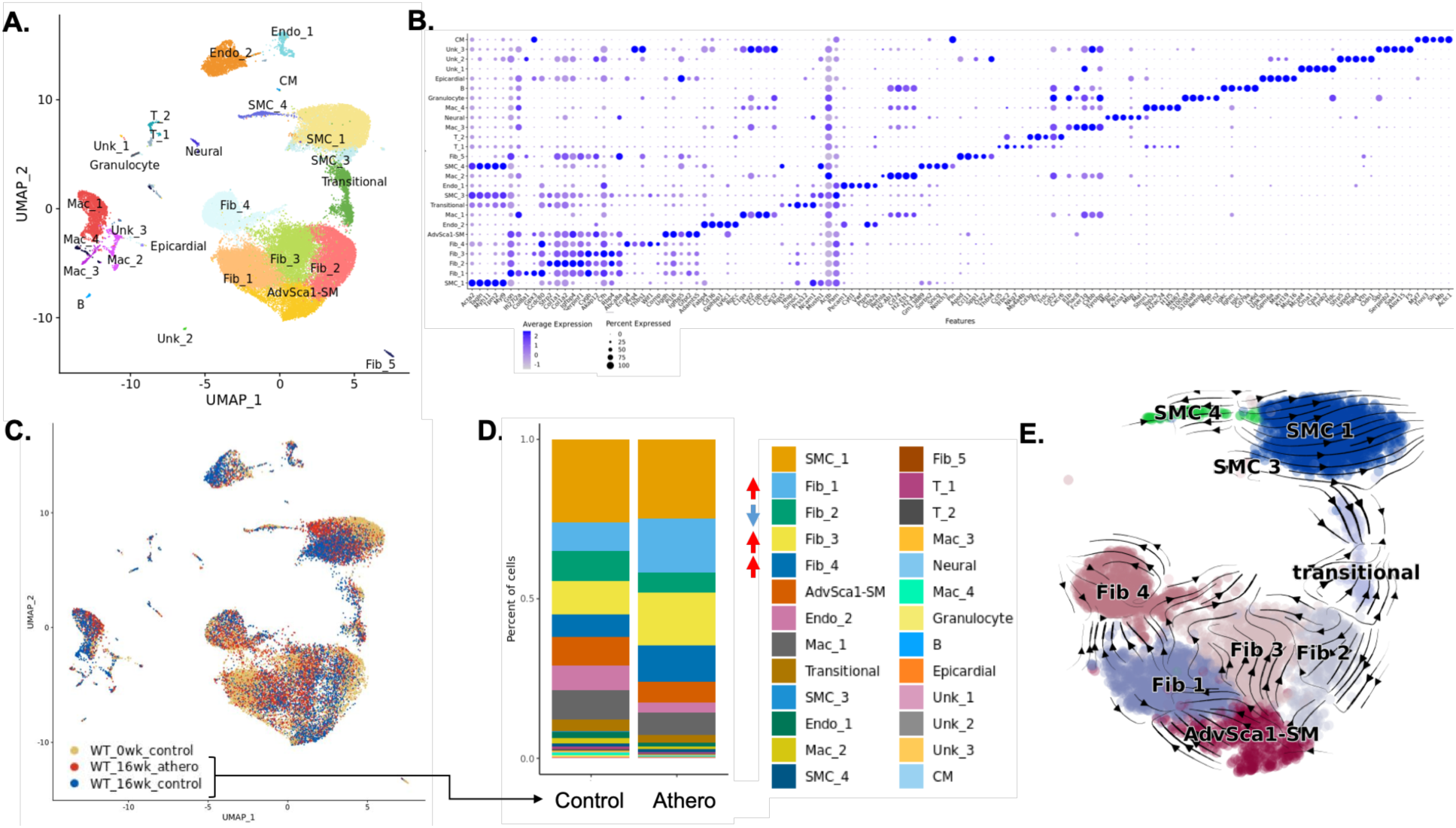
Single cell RNA sequencing analysis demonstrates major shifts in vascular cell types in the setting of atherosclerosis. The aortic sinus, aortic arch, brachiocephalic artery, and carotid arteries were harvested at baseline and after 16 weeks of normal high fat chow and processed for scRNA-Seq. **A.** UMAP projection of all cells that passed quality control. Cluster identity was assigned using representative gene expression profiles. **B.** Dot plot showing the top 5 unique genes that define each cluster. **C.** UMAP of all cells from baseline, 16-week control, and 16-week atherogenic samples. **D.** Stacked bar plot of YFP^+^ and YFP^-^ cells from 16-week control and atherogenic samples. Arrows indicate increases (red) or decreases (blue) in the cell population as a result of atherosclerosis. **E.** RNA velocity analysis of all cells after 16 weeks of atherogenic diet.

### AdvSca1-SM cells preferentially differentiate into fibroblasts or remain in a stem-like state in atherosclerosis, with minor contribution to SMC populations

Leveraging our AdvSca1-SM lineage tracing mouse model, we investigated specific changes in the YFP^+^ AdvSca1-SM and AdvSca1-SM-derived cell populations in the setting of atherosclerosis. YFP^+^ cells were primarily found in the stemlike AdvSca1-SM, and fibroblast-like Fib_1, Fib_2, and Fib_3 clusters (Figure 3A). These clusters expressed high levels of myofibroblast (*Tcf21*) and extracellular matrix markers (*Col1a1, Col1a2, Col3a1, Lum, Dcn*) and stem cell markers (*Ly6a, Cd34, Scara5, Pi16*)(Figure 3B). However, the distribution of these markers varied between the clusters, with the AdvSca1-SM cell representing the most stem-like phenotype and Fib_2 having elevated collagen gene expression. Aortic roots stained for YFP and Sca1 exhibited a large population of YFP^+^/Sca1^+^ cells in the adventitia (Figure 3C). Flow cytometry analysis of animals after 16 weeks of atherogenic diet supported these findings, demonstrating a considerable proportion of YFP^+^/Sca1^+^ stemlike AdvSca1-SM cells (Figure 3D). It should be pointed out that scRNA-Seq and flow were performed on whole arterial tissue compared to immunofluorescence analysis of plaque regions. As atherosclerosis is a focal disease, many of these stemlike AdvSca1-SM cells are likely from regions of uninvolved tissue. We confirmed expression of stem cell and fibroblast markers in YFP^+^ cells using immunofluorescence and RNAscope microscopy Figure 3C-E). YFP^+^ cells in both the adventitia and plaque were found to express the fibroblast marker, lumican (Lum). Although the primary route of AdvSca1-SM cell differentiation was towards a fibroblast phenotype or remaining as a stem cell, a subset YFP^+^ AdvSca1-SM cells differentiated towards a SMC fate. YFP^+^ cells found in the SMC clusters were found to express genes specific to mature vascular SMCs, including *Acta2, Myh11, Cnn1*, and *Tagln* (Figure 3F). This differentiation profile was confirmed using immunofluorescence microscopy, in which we identified YFP^+^/αSMA^+^ cells comprising the plaque fibrous cap (Figure 3G). Finally, we identified extremely rare differentiation of AdvSca1-SM cells into other cell types, including macrophages and endothelial cells (Extended Data Figure 6).

**Figure 3.**
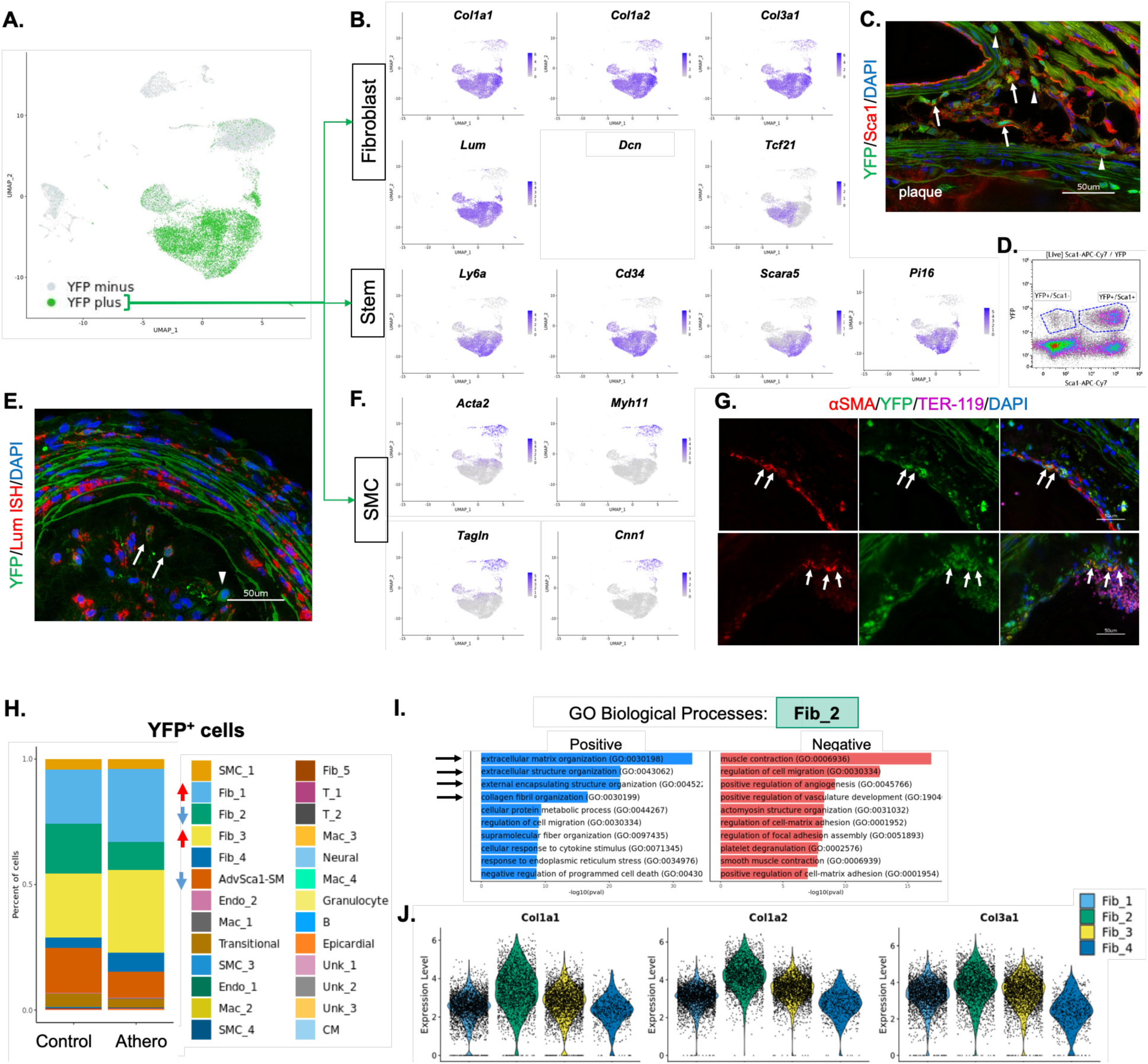
AdvSca1-SM cells primarily differentiate into fibroblasts or remain in a stem-like state in atherosclerosis, but can also differentiate into SMCs. **A.** Feature plots showing the distribution of YFP^+^ and YFP^-^ cells on the UMAP. **B.** Feature plots of major fibroblast (*Col1a1, Col1a2, Col3a1, Dcn, Lum*, and *Tcf21*) and stem cell markers (*Ly6a*/Sca1, *Cd34*, *Scara5*, *Pi16*) in YFP^+^ cells at all time points. **C.** Representative aortic root image from 24-week plaque stained for YFP (green), Sca1 (red), and DAPI (blue). Arrows indicate YFP^+^/Sca1^+^ cells in the adventitia; arrowheads indicate YFP^+^/Sca1^-^ cells. * = cardiomyocyte autofluorescence. **D.** Aortic sinus, aortic arch, brachiocephalic artery, and carotid arteries from 16-week control or atherosclerotic mice were processed for FACS analysis. Representative image of a YFP and Sca1 density plot from an atherogenic animal. **E.** Double RNAscope and immunofluorescence image of an aortic root from 24-week atherogenic animals showing lumican (*Lum*; fibroblast cell marker; red) mRNA and YFP (green). Arrows indicate YFP^+^/lumican^+^ cells; arrowheads indicate YFP^+^/lumican^-^ cells. **F.** Feature plots of SMC genes (*Acta2, Cnn1, Mhy11, Tagln*) in YFP^+^ cells at all time points. **G.** 24-week aortic root plaques were stained for YFP (green), αSMA (red), Ter-119 (erythroid cells; magenta), and DAPI (blue). Arrows indicate YFP^+^/αSMA^+^ cells forming the fibrous cap of the plaque. **H.** Stacked bar plot showing phenotypic shifts in YFP^+^ cells between 16-week control and atherogenic samples. Arrows indicate increases (red) or decreases (blue) in the cell population as a result of atherosclerosis. **I.** GO Biological Process between the Fib_2 cluster and all other cell clusters. Arrows indicate the top 4 processes positively associated with Fib_2. **J.** Violin plots demonstrating the stronger collagen/extracellular matrix gene signature (*Col1a1, Col1a2*, and *Col3a1*) in Fib_2 compared to Fib_3. All scale bars, 50μm.

As expected, the proportion of YFP^+^ cells in each of the cell clusters was altered in mice on atherogenic diet compared to controls. Specifically, there were increases in YFP^+^ cells in Fib_1 and Fib_3 clusters with a corresponding decrease in Fib_2 and AdvSca1-SM cell clusters in the setting of atherosclerosis (Figure 3H). To elucidate the functional importance of these cell shifts, we further characterized the phenotypes of the different fibroblast clusters using GO Biological Process pathway analysis. We determined that compared to all other cell clusters, Fib_2 was enriched for pathways related to extracellular matrix (ECM) and collagen organization (Figure 3I). These findings were confirmed by comparing average expression of collagen genes between the various fibroblast clusters (Figure 3J). Collectively, these data support that chronic hyperlipidemia-induced atherosclerosis promotes activation and a shift in the phenotype of AdvSca1-SM cells predominantly to myofibroblast-like cells, but also toward a mature SMC phenotype. These findings are consistent with immunofluorescence data demonstrating significant contributions of AdvSca1-SM cells to adventitial remodeling, medial repair, and plaque formation (both fibrous cap and core plaque cells).

### Dynamic exchange between AdvSca1-SM cells and SMCs in the setting of atherosclerosis

RNA velocity analysis of our data (Figure 1E) revealed that not only do AdvSca1-SM cells differentiate away from their most stem-like state and into other cell types, but also that there are a variety of cell transitions occurring in the fibroblast and SMC clusters. Of particular interest was the Transitional cluster which occupies UMAP space between SMCs and AdvSca1-SM/fibroblast clusters and shows a bidirectional differentiation trajectory. The first trajectory demonstrated a shift from AdvSca1-SM/fibroblast clusters toward a SMC phenotype, a trajectory which we confirmed (Figures 3F&G). However, we also observed a reverse trajectory from non-AdvSca1-SM-derived mature SMCs to AdvSca1-SM/fibroblast-like clusters. Further characterization of this Transitional cluster revealed two unique subpopulations, an upper cluster which expresses more SMC genes and is largely YFP^-^, and a lower cluster, which expresses more stem/fibroblast genes with the majority of cells being YFP^+^ (Figure 4A). Further supporting the identity as an intermediate or transitional cell population, KEGG pathway analysis of the Transitional cluster showed it to be less contractile than mature SMC clusters, but more contractile than the fibroblast/AdvSca1-SM clusters (Figure 4B). These data support differentiation of YFP^+^ AdvSca1-SM cells toward a SMC phenotype, but also YFP^-^ mature SMC reprogramming toward a stemlike AdvSca1-SM phenotype, consistent with our original discovery of SMC-derived AdvSca1-SM cells^14^. To confirm this, we exposed SMC lineage tracing mice (Myh11-Cre^ERT^/Rosa26-YFP) to tamoxifen to label SMCs, then put them on the same atherogenic regimen as the AdvSca1-SM lineage tracing mice. We found that SMC-derived YFP^+^ cells were found in the vascular adventitia where they co-expressed Sca1, indicating the transition of a mature, contractile SMC into multipotent progenitor cells (Figure 4C). Interestingly, we also observed YFP^-^ cells (i.e. non-SMC-derived) within the media of the vessel (Figure 4C). These data demonstrate the dynamic nature of these cells and support a novel mechanism whereby SMCs maintain and replenish the resident AdvSca1-SM stem cell pool while SMC-derived AdvSca1-SM cells redifferentiate toward a SMC fate to contribute to vessel repair/homeostasis.

**Figure 4.**
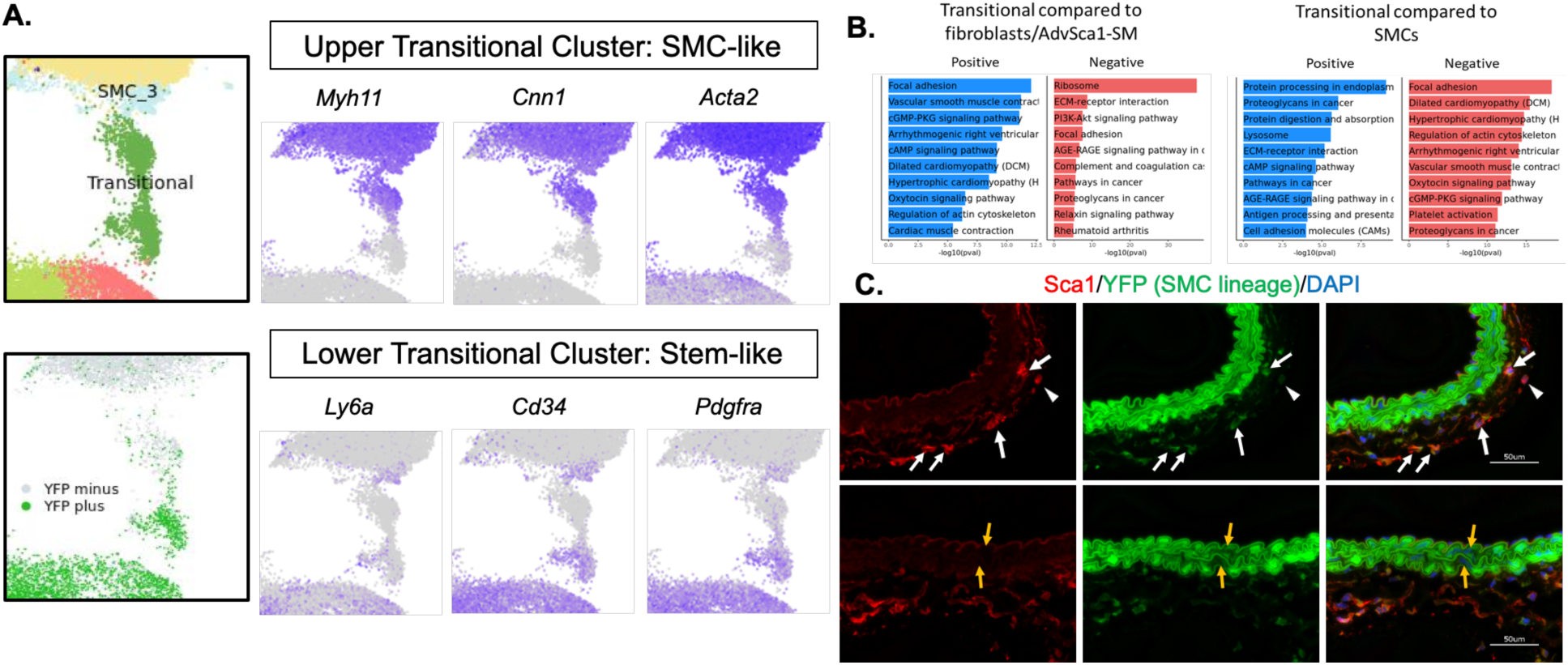
Dynamic, bidirectional differentiation between AdvSca1-SM cells and SMCs in the setting of atherosclerosis. **A.** The Transitional cluster consists of YFP^+^ and YFP^-^ cells and exhibits features of both SMCs and AdvSCa1-SM cells, roughly divided in half (bottom). Feature plots show that the upper portion of the transitional cluster expresses more contractile SMC genes (*Acta2, Myh11, Cnn1*), whereas the lower portion expresses more mesenchymal stem cell markers (*Ly6a, Cd34, Pdgfra*). **B.** KEGG Pathway differences between the Transitional cluster and the AdvSca1-SM/fibroblast clusters or SMC clusters. **C.** Representative image of 14-week atherogenic braciocephalic artery from SMC lineage tracing mice (Myh11-CreERT^+/-^/Rosa26-YFP^+/+^) stained for YFP (green; expression indicates SMCs, not AdvSca1-SM). Arrows indicate adventitial YFP^+^/Sca1^+^ cells; arrowheads indicate adventitial progenitors not expressing YFP (YFP^-^/Sca1^+^); orange arrows indicate DAPI positive cells in the vessel media that are negative for YFP.

### Depletion of stemness transcription factor KLF4 specifically in AdvSca1-SM cells alters their differentiation trajectory in atherosclerosis

Having established the contribution of AdvSca1-SM cells to atherosclerosis progression, we sought to determine the effect of genetic modulation of these cells on plaque progression and complexity. Given the integral function of KLF4 in the SMC-to-AdvSca1-SM cell transition and maintenance of the progenitor phenotype, KLF4 was selectively depleted in AdvSca1-SM cells as we previously published^14, 15^. Following tamoxifen treatment to induce YFP reporter knock-in and KLF4 knockout (in Klf4^fl/fl^ mice), mice were put on the same atherogenic regimen as previously described (Figure 1A). No differences were observed between WT and Klf4 KO mice in baseline weight or weight change and total cholesterol levels after 24 weeks of atherogenic conditions (Extended Data 5). Arteries from WT and Klf4 KO mice were harvested for scRNA-Seq and histological analyses. After filtering sequencing data for AdvSca1-SM cell-derived YFP^+^ cells following 16 weeks of atherogenic treatment, we observed significant shifts in cell populations as a consequence of KLF4 depletion in AdvSca1-SM cells (Figure 5A). In particular, we observed an increase in Fib_2 with a corresponding decrease in Fib_3 in arteries from the Klf4 KO mice compared to WT, precisely the opposite shift observed when comparing WT animals on control diet to atherogenic diet (Figure 5B). This increase in Fib_2 indicates that AdvSca1-SM cell-derived fibroblasts in Klf4 KO mice transition to a phenotype characterized by enriched ECM and collagen deposition. Additionally, comparing Fib_2 and Fib_3 populations between WT and Klf4 KO mice, cell populations from Klf4 KO mice expressed higher levels of collagen genes (*Col1a1* and *Col1a2)*, further supporting a shift in phenotype of Klf4 KO AdvSca1-SM cell-derived YFP^+^ fibroblasts towards a collagen deposition phenotype that potentially contributes to a protective fibrous cap (Figure 5C). Finally, there was a shift in the phenotype of AdvSca1-SM-derived Transitional cells toward the mature SMC phenotype (Figure 5D), supporting a role in medial repair.

**Figure 5.**
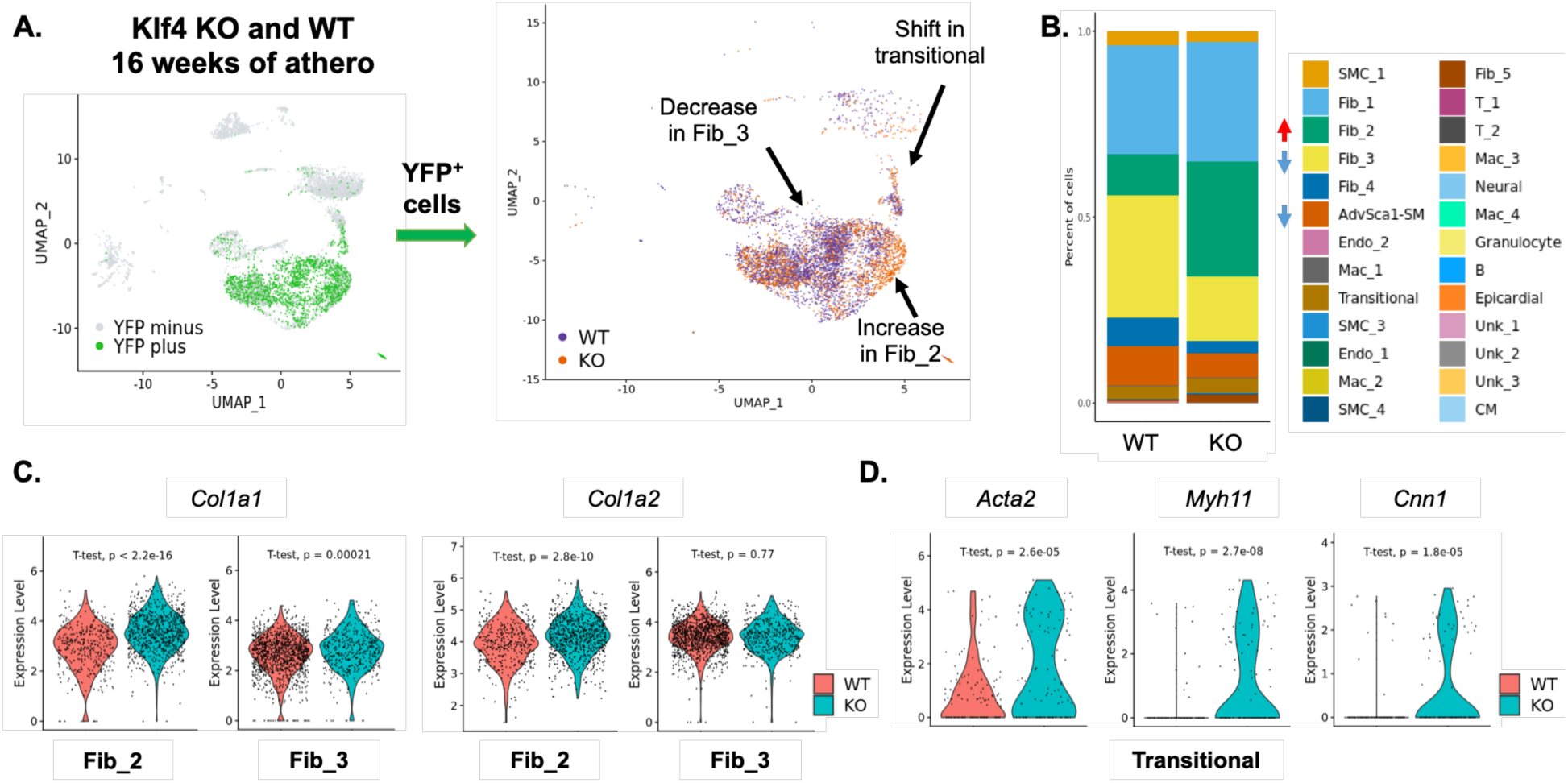
Depletion of KLF4 in AdvSca1-SM cells modifies the fate of YFP^+^ cells in the setting of atherosclerosis. Arteries from 16-week WT and Klf4 KO AdvSca1-SM lineage atherogenic mice were processed for scRNA-Seq. **A.** [**Left**] UMAP projection of YFP^+^ and YFP^-^ cells from WT and Klf4 KO AdvSca1-SM cell lineage tracing mice. [**Right**] UMAP projection showing the different fates of YFP^+^ cells in atherosclerosis as a consequence of depleting KLF4 in AdvSca1-SM cells. Arrows indicate major phenotypic shifts in KO mice. **B.** Stacked bar graph of YFP^+^ cells from WT and Klf4 KO mice after 16-week atherogenic diet. Arrows indicate increases (red) or decreases (blue) in cell populations as a function of Klf4 depletion. **C.** Violin plots showing higher expression of ECM related genes (*Col1a1, Col1a2*) in YFP^+^ Fib_2 cluster from Klf4 KO mice compared to WT mice after 16 weeks of atherogenic diet. *Col1a1* is also higher in YFP^+^ Fib_3 cluster cells from Klf4 KO mice compared to WT mice, but there is no difference with *Col1a2* in this cluster. **D.** Violin plots showing higher expression of SMC contractile genes (*Acta2, Myh11, Cnn1*) in YFP^+^ Transitional cluster cells from Klf4 KO mice compared to WT mice after 16 weeks of atherogenic diet.

### Phenotypic modulation of AdvSca1-SM cells via KLF4 depletion modifies the fate of non-AdvSca1-SM-derived cells

While we anticipated that KO of KLF4 specifically in AdvSca1-SM cells would alter their differentiation patterns during atherosclerosis progression, we also observed significant shifts in the phenotype of YFP^-^ non-AdvSca1-SM-derived cell populations in response to AdvSca1-SM cell Klf4 KO (Figure 6A). The most notable changes were in the gene expression profile of the major SMC cluster (SMC_1) as well as an overall decrease in the primary macrophage cluster (Mac_1 cluster)(Figure 6B). Further examination of the shift within the SMC_1 cluster revealed that SMCs from Klf4 KO mice exhibited slightly elevated expression of smooth muscle contractile genes (Figure 6C). Investigation of Mac_1 revealed that macrophages from Klf4 KO mice expressed lower levels of *Ccl2* (MCP-1), suggesting a reduced inflammatory phenotype and diminished recruitment of additional macrophages to the lesion (Figure 6D). Flow cytometry analysis of arteries from both WT and Klf4 KO mice after 24 weeks of atherogenic diet revealed a non-significant trend toward a decrease in F4/80^+^ macrophages in the whole vascular digest (Figure 6E). As it has been well established that macrophages play an essential role in atherosclerosis progression, we sought to further define the phenotypic changes occurring in Mac_1 in the KO animals. Since Klf4 was selectively depleted only in AdvSca1-SM cells and Mac_1 population is almost entirely YFP^-^ non AdvSca1-SM cell-derived, we hypothesized a paracrine signaling axis resulted in the phenotypic shifts observed in Mac_1 macrophages from Klf4 KO mice. We investigated this possibility using Cell Chat, which infers cell-cell communications in scRNA-Seq datasets. We identified a high likelihood of fibroblast/AdvSca1-SM cell collagen-to-Mac_1 SDC4 or CD44 signaling. Additionally, these signaling pathways were predicted to increase in KO animals (Figure 6F). Previous research has indicated that low levels of SDC4 in macrophages promotes atherosclerosis by polarizing macrophages towards a more pro-inflammatory state^24^. This polarization was shown to occur through SDC4-mediated changes in ABCA1 and ABCG1 signaling, the ATP-binding cassette transporters essential for cholesterol efflux to HDL. Interestingly, we found that, compared to WT mice, macrophages from Klf4 KO mice expressed higher levels of *Sdc4* and *Abcg1,* but not *Abca1* (Figure 6G). These findings highly support the concept that not only are macrophages from WT animals more likely to recruit other macrophages via MCP-1, but they are also likely to be more pro-inflammatory and pro-atherogenic. Collectively, our data support a model in which manipulation of the phenotype of AdvSca1-SM cells results in autonomous effects on AdvSca1-SM cell differentiation and function as well as non-autonomous atheroprotective effects on additional cells of the atherosclerotic microenvironment.

**Figure 6.**
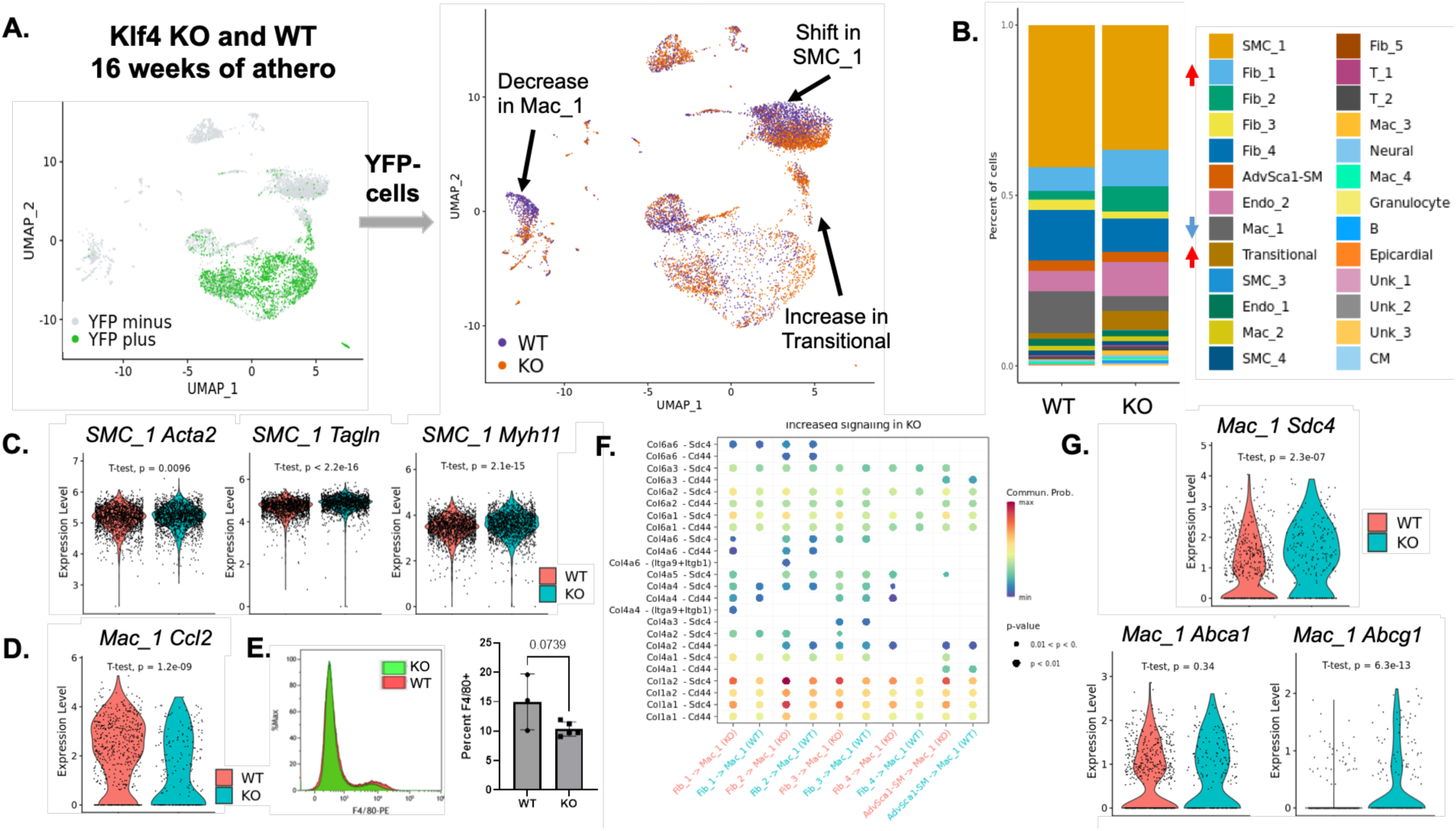
KLF4 depletion in AdvSca1-SM cells alters the fate of YFP^-^ non-AdvSca1-SM-derived cells. Arteries from 16-week WT and Klf4 KO AdvSca1-SM lineage atherogenic mice were processed for scRNA-Seq. **A.** [**Left**] UMAP projection of YFP^+^ and YFP^-^ cells from WT and Klf4 KO AdvSca1-SM cell lineage tracing mice. [**Right**] UMAP projection showing the different fates of YFP^-^ cells in atherosclerosis as a consequence of depleting KLF4 in AdvSca1-SM cells. Arrows indicate major phenotypic shifts in KO mice, including changes to the SMC and macrophage clusters. **B.** Stacked bar graph of YFP^-^ cells from WT and Klf4 KO mice after 16-week atherogenic diet. Arrows indicate increases (red) or decreases (blue) in the cell population as a result of KLF4 depletion. **C.** Violin plot showing expression of SMC contractile genes (*Acta2, Tagln, Myh11*) in YFP^-^ cells in the major SMC cluster from Klf4 KO mice compared to WT mice. **D.** Violin plot showing expression of *Ccl2* in YFP^-^ cells of the major macrophage cluster from Klf4 KO mice compared to WT mice. **E.** Flow cytometry analysis of single cell arterial digests from both WT and Klf4 KO mice after 24 weeks of atherogenic diet. **F.** CellChat analysis was performed on the fibroblast/AdvSca1-SM and Mac_1 clusters. Bubble plot showing elevated levels of COL1A1/COL1A2 from the fibroblast clusters signaling to SDC4 in Mac_1 in the setting of atherosclerosis. **G.** Violin plots show expression of *Sdc4* and *Abcg1* in Mac_1 from Klf4 KO mice in the setting of atherosclerosis. *Abca1* is not significantly different between the genotypes.

### Klf4 knock out in AdvSca1-SM cells does not reduce plaque burden, but promotes a more stable plaque phenotype

To ascertain the physiological consequences of the AdvSca1-SM cell-specific Klf4 KO, we examined overall plaque burden. In the aortic root, aortic arch, and full vascular tree, there were no significant differences in total plaque burden between the WT and Klf4 KO mice at either early or late time points (Figures 7A and 7B, Extended Data Figure 7). However, there was a significant reduction in total necrotic core area in the plaques from Klf4 KO mice compared to WT mice, indicative of a more stable plaque (Figure 7B). Additionally, plaques from Klf4 KO mice showed fewer cholesterol clefts within the plaques (Figure 7C), supporting our earlier findings suggesting impaired cholesterol efflux to HDL in Mac_1 from WT compared to Klf4 KO mice. Quantification of αSMA^+^ fibrous cap area in the aortic root revealed increased fibrous cap thickness in plaques from the Klf4 KO mice (Figure 7D). Our scRNA-Seq and flow data revealed a non-significant trend toward an overall reduction in macrophages from Klf4 KO mice, but these approaches involved analysis of both diseased and non-diseased regions of the vessel (Figure 6E). Since atherosclerosis is a focal disease, we also used an immunofluorescence approach to examine macrophage accumulation specifically in plaque regions. Compared to WT mice, we found significantly reduced accumulation of CD68^+^ macrophages in the plaques from the Klf4 KO mice (Figure 7E). Finally, we investigated the plaque structure using Masson’s Trichrome, which demonstrated increased collagen deposition in plaques from Klf4 KO mice compared to WT controls, supporting the shifts in fibroblast populations we identified in the scRNA-Seq experiments (Figure 7F). Collectively, these data point to a more stable plaque phenotype in mice with AdvSca1-SM-specific Klf4 KO compared to WT controls.

**Figure 7.**
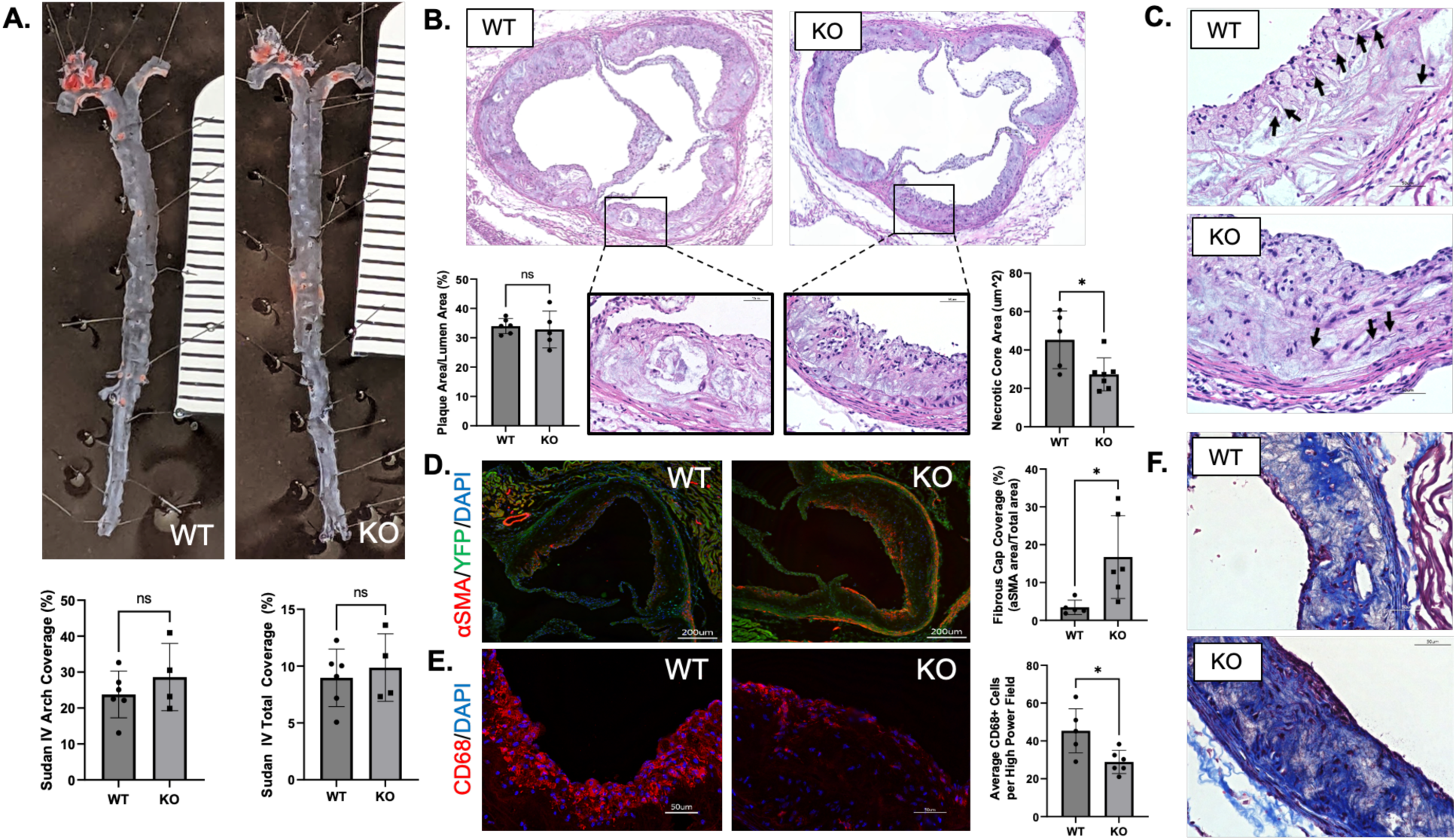
Depletion of KLF4 in AdvSca1-SM cells doesn’t alter overall plaque burden, but reduces late-stage plaque complexity. **A.** The aortic arch and descending aorta from 24-week atherogenic WT and Klf4 KO mice were stained using Sudan IV to assess overall atherosclerotic plaque burden. Quantification of staining is shown in the lower graphs. **B.** Representative H&E images of aortic root cross sections from WT and Klf4 KO mice. Insets show higher magnification of the plaque structure with significantly smaller necrotic cores in Klf4 KO mice compared to WT mice. Quantification of necrotic cores is shown in the lower right graph. **C.** Representative H&E images of stained aortic root cross sections from WT and Klf4 KO mice. Arrows indicate cholesterol clefts. **D.** Representative immunofluorescent images of aortic root cross sections from WT and Klf4 KO mice for YFP (green), αSMA (red), and DAPI (blue). Quantification of staining is shown to the right. **E.** Representative immunofluorescent images of aortic root cross sections from WT and Klf4 KO mice for CD68 (red) and DAPI (blue). Quantification of staining is shown to the right. **F.** Representative Masson’s Trichrome images from WT and Klf4 KO mice aortic roots with collagen (blue), cytoplasm (pink), and cell nuclei (brown). Scale bars for all images are 50 μm, except for panel D (200 μm).

## DISCUSSION

In this study, we established the importance of an under-studied cell population, AdvSca1-SM stem cells, to atherosclerosis progression. Consistent with our previous research in the setting of acute vascular injury, this is the first report demonstrating that AdvSca1-SM cells and AdvSca1-SM-derived cells are found throughout the vessel wall contributing to pathological atherosclerotic lesions and adventitial remodeling. We also established the major differentiation pathways of AdvSca1-SM cells to fibroblasts and SMCs, with minor contributions to other cell populations, including endothelial cells, macrophages, and adipocytes. AdvSca1-SM specific genetic depletion of the reprogramming-associated transcription factor, KLF4, drastically altered the differentiation trajectories of AdvSca1-SM cells as well as the phenotypes of other vascular cell types, including macrophages and SMCs. Finally, we found that these autonomous and non-autonomous effects of KLF4 knock out resulted in atheroprotection through increased plaque stability, as measured by decreased necrotic core size, increased fibrous cap thickness, and decreased macrophage accumulation (Figure 8).

**Figure 8.**
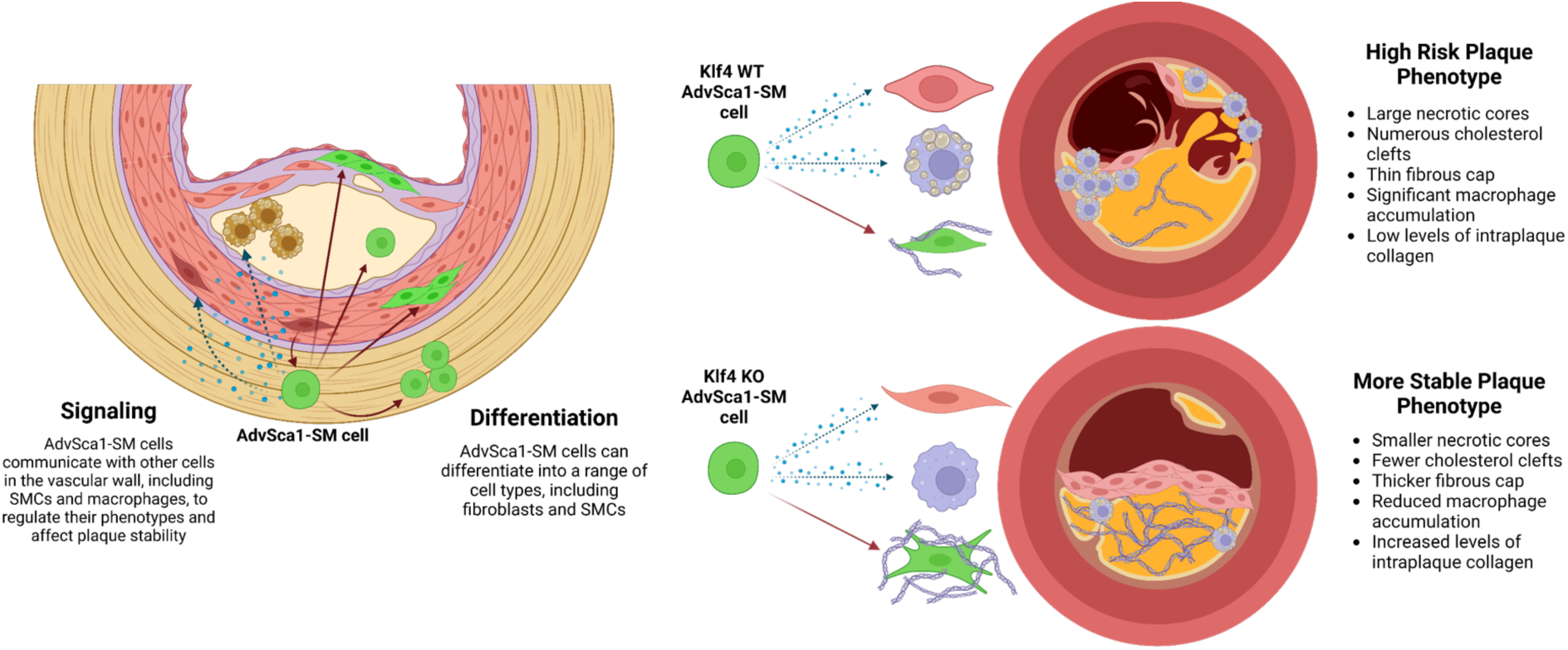
Cartoon describing the multifaceted roles of AdvSca1-SM cells in atherosclerosis and their ability to alter plaque phenotype. In the setting of atherosclerosis, AdvSca1-SM cells are capable of differentiating into a variety of cell types, predominantly fibroblasts and SMCs, and migrating throughout the vascular wall and atherosclerotic plaque. In addition to directly differentiating into other cell types, AdvSca1-SM cells signal to other cells, including macrophages, fibroblasts, and SMCs, to alter their phenotypes. Depletion of the stemness-associated transcription factor, KLF4, selectively in AdvSca1-SM cells results in altered AdvSca1-SM-derived cell, macrophage, and SMC phenotypes, leading to a more stable plaque phenotype.

Previous studies of the adventitia in the setting of atherosclerosis have focused on the role of the vasa vasorum (VV) in regulating atherosclerosis via the “outside-in hypothesis”^6–9^. Research has highlighted the importance of these microvessels by showing that neovascularization is correlated with plaque size while inhibition of plaque neovascularization reduced atherosclerosis progression^8, 25–28^. In this study, we identified YFP^+^ AdvSca1-SM-derived cells surrounding adventitial microvessels, suggesting a role for AdvSca1-SM cells in VV expansion and potentially plaque neovascularization. However, it is important to note that the majority of AdvSca1-SM cells were not found associated with the microvessels, but nonetheless greatly influenced plaque complexity through contribution to plaque core cells as well as fibrous cap cells. These findings support the importance of resident SMC-derived adventitial AdvSca1-SM stem cells in atherosclerosis.

Our finding that the primary differentiation trajectory for AdvSca1-SM cells in the setting of atherosclerosis is towards a fibroblast phenotype closely mirrors our previous findings in the setting of acute vascular injury^15, 16^. Additionally, other groups using similar mouse models have indicated that these adventitial progenitor cells are involved in vascular calcification and neoinima formation in the setting of chronic kidney disease and in the pathological profibrotic response in multiple organ systems^29, 30^. However, in contrast to acute injury models in which perivascular fibrosis is detrimental, there is an important function for modulated SMC- and fibroblast-associated collagen deposition in protecting against necrotic core formation and plaque rupture^31^. Importantly, our findings of increased collagen I and III expression by AdvSca1-SM-derived fibroblasts, as observed in AdvSca1-SM Klf4 KO mice compared to WT mice, supports the concept that manipulating the phenotype of AdvSca1-SM cells confers a more stable/protective plaque phenotype, thereby making targeting of these cells an attractive therapeutic target.

A major focus of this study was the effect of *Klf4* depletion on AdvSca1-SM cell phenotype and function as well as on the plaque microenvironment. KLF4 has been widely recognized as one of the key transcription factors necessary for induction of pluripotent stem cells (iPSCs)^32^. In the vasculature, we demonstrated that KLF4 induction is essential for SMC reprogramming and maintenance of the AdvSca1-SM stem cell phenotype. AdvSca1-SM-specific KLF4 depletion results in spontaneous differentiation toward a myofibroblast pro-fibrotic phenotype^14, 15^. Others have established that KLF4 is a potent repressor of the mature contractile phenotype and that KLF4 depletion in SMCs results in a reduction in atherosclerotic plaque burden as well as aortic aneurysm formation^33, 34^. Adding translational significance was the identification of *Klf4* by genome-wide association studies (GWAS) in human populations as a coronary artery disease risk locus^23, 35^. However, whether KLF4 is protective or deleterious seems to be context dependent. Contrary to the previous findings in SMCs, KLF4 depletion in endothelial and myeloid cells drives atherosclerosis progression^36, 37^. In this study, we established that KLF4 in AdvSca1-SM cells is deleterious and that depletion results in more stable atherosclerotic plaques. This finding supports much of the prior research in vascular SMCs, but also serves to highlight the importance of elucidating the cell-specific effects of KLF4 in different disease processes.

One of the most unexpected findings was the ability of AdvSca1-SM cells to regulate the phenotype of plaque-associated macrophages. Our data suggest that cross communication may be a result of collagen produced by AdvSca1-SM cells and AdvSca1-SM-derived fibroblasts acting as a ligand for SDC4 on the surface of macrophages. A recent study from Hu et al. demonstrated that macrophage surface SDC4 is decreased in the setting of atherosclerosis, and that this reduction in SDC4 is both proinflammatory and pro-atherogenic^24^. It is believed that the proinflammatory macrophage phenotype is due to a decrease in ABCA1/ABCG1, which act as the rate limiting steps for cholesterol efflux and reverse cholesterol transport^38, 39^. Further highlighting the importance of the ABCA1/ABCG1 axis are studies showing that macrophage-specific deficiency of ABCA1 and ABCG1 promotes plaque inflammation and signals to bone marrow progenitors to produce more monocytes; it has also been demonstrated that ABCA1 and ABCG1 are major suppressors of plaque associated leukocytosis, which confers an atheroprotective effect^40, 41^. These studies emphasize the central role of macrophage cholesterol efflux in the regulation of atherosclerosis. Importantly, our data with Klf4 KO mice compared to WT mice suggests that loss of Klf4 in AdvSca1-SM cells results in enhanced collagen production and increased *Sdc4/Aba1/Abcg1* expression with a corresponding reduction in *Ccl2* expression, which likely contributes to the observed plaque protective effects.

One notable limitation in studying AdvSca1-SM cells is the dynamic nature of this cell population. AdvSca1-SM cells are poised to respond to changes in the vascular environment, and as such are capable of quickly differentiating away from their stemlike state. Similarly, SMC reprogramming to AdvSca1-SM cells is a dynamic process. Using our lineage tracing system, we were able to track these cells over time even as they underwent major cell transitions. However, due to the dynamic SMCs-to-AdvSca1-SM cell transition, as shown in both our previous work and this study^14^, newly generated AdvSca1-SM cells will be formed, but will not be labeled with the AdvSca1-SM reporter, nor will KLF4 be depleted in the case of Klf4 KO mice. This cell turnover is likely to have diluted some of the findings in this study and future investigation into the timing and frequency of these reprogramming events is likely warranted. Despite these technical concerns, the dynamic nature of the ongoing reciprocal SMC-to-AdvSca1-SM transitions identified by RNA velocity analysis suggest that understanding the mechanisms involved in these transitions might provide sensitive therapeutic targets.

To summarize, in this study we used a combination of histological and single cell sequencing approaches to define the multifaceted contributions of AdvSca1-SM cells to atherosclerosis, both through direct differentiation into other cell types and through signaling to SMCs and macrophages. Additionally, we revealed the potential for modulation of these resident vascular progenitor cells to regulate atherosclerotic plaque stability. Collectively this work emphasizes the continuing importance of research into the vascular adventitia as a driver of atherosclerosis and a potential therapeutic target.

## MATERIALS AND METHODS

### RESOURCE AVAILABILITY

#### Lead contact

Further information and requests for resources and reagents should be directed to and will be fulfilled by the lead contact, Mary C.M. Weiser-Evans.

#### Materials Availability

The materials generated from this work may be shared with the scientific community after appropriate invention disclosure(s) and adequate protection has been achieved by the University of Colorado’s Technology Transfer Office. Samples of research materials will be made available to the research community following completion of the University of Colorado’s Material Transfer Agreement.

#### Data and Code Availability

This research generated data from experiments using genetic mouse models. Datasets for single cell RNA-Seq have been deposited in the NCBI Gene Expression Omnibus site (https://www.ncbi.nlm.nih.gov/geo/). scRNA-Seq data was analyzed with using Seurat and other R packages; code for this analysis has been deposited in the Weiser-Evans Lab GitHub (https://github.com/weiserevanslab/adubner).

### EXPERIMENTAL MODEL AND SUBJECT DETAILS

#### Mice

All mice for this project were approved by our Institution IACUC, protocol #00066 Weiser-Evans. Animal numbers were approved by the University of Colorado Anschutz Medical Campus Animal Institute Committee.

This research utilized a variety of transgenic mice generated from widely available strains. Gli1-Cre^ERT^ (JAX 007913), Myh11-Cre^ERT^ (Stephan Offermanns, Max-Planck-Institute for Heart and Lung Research, Germany; now available at JAX 019079), and Rosa26-YFP (JAX 006148) mice were all obtained from The Jackson Laboratory. Klf4 floxed mice were obtained from Dr. Klaus H. Kaestner (University of Pennsylvania, Philadelphia). All mice currently reside as colonies in Dr. Weiser-Evans’ laboratory.

Lines generated by breeding:

a). Gli1-Cre^ERT^/Rosa26-YFP (AdvSca1-SM lineage tracing mouse, a.k.a. WT)
b). Gli1-Cre^ERT^/Rosa26-YFP/Klf4^flox/flox^ (AdvSca1-SM lineage tracing mouse with AdvSca1-SM specific deletion of Klf4, a.k.a. Klf4 KO)
c). Myh11-Cre^ERT^/Rosa26-YFP (SMC lineage tracing mouse)

All mice bred to Gli1-Cre^ERT^ transgenic mice are inducible systems for AdvSca1-SM reporter knock-in and/or AdvSca1-SM-specific Klf4 knock out; mice bred to Myh11-Cre^ERT^ are inducible systems for SMC reporter knock-in. Experimental mice were maintained heterozygous for the Gli1-Cre^ERT^ or hemizygous for Myh11-Cre^ERT^ and homozygous for YFP reporter and Klf4 flox. Age-matched male and female mice were used for all AdvSca1-SM lineage mouse experiments, with tamoxifen injections beginning between 6 and 8 weeks of age, and initiation of atherogenic or control treatment regimens beginning between 8 and 10 weeks of age. Only male mice were used for the SMC lineage tracing mouse experiments as the Cre was inserted on the Y chromosome. Weight was measured at the beginning and end of the studies to identify any outliers. Starting weight was not significantly different between WT and KO mice at baseline, nor was there a difference between the genotypes in weight change during the experiments (Extended Data Figure 5).

Experimental mice were housed in the AALAC accredited RC-2 vivarium at the University of Colorado Anschutz Medical Campus. The vivarium is maintained at standard sub-thermoneutral temperatures (22-26°C) and with a 12-hour light-dark cycle. All animals had *ad libitum* access to both food and water. Atherogenic mice received Envigo TD.02028 (42.6% fat, 1.3% cholesterol, and 0.5% cholic acid) or Research Diets D05060402 (42.8% fat, 1.5% cholesterol, and 0.5% cholic acid), whereas control animals received the standard diet Envigo 2920x (16% fat); all diets were irradiated prior to transfer into the animal facility. Animal health was monitored daily by veterinary staff, and animals were humanely euthanized per protocol in the event of any significant illness or injury. Cages were changed every 2 weeks and contained enrichment materials, including nesting material (e.g. Nestlet, brown shredded paper, paper towels, etc.) and/or cage furniture. Animals on atherogenic diet received front and rear nail trims as needed to minimize scratch trauma resulting from diet-induced dermatitis.

### METHOD DETAILS

#### Mouse Genotyping

All mice used in this project were bred and maintained by our lab in the University of Colorado RC-2 Vivarium. At the time of weaning, mice were ear tagged and ear snipped for genotyping and future identification. Ear snips were then processed using Extracta DNA Prep for PCR-Tissue (QuantaBio, 95091-025) per manufacturer protocol. DNA samples were stored at -20°C until ready for genotyping.

Prior to use in experiments, mice were genotyped using the recommended primers and PCR programs from The Jackson Laboratory. For each primer set, a master mix was prepared with 10 μl AccuStart II PCR SuperMix (QuantaBio 95137), 1 μl of each primer (listed below, diluted 1:10 in molecular grade water for use), and molecular grade water to bring reaction total up to 17 μl per sample. Master mix was loaded into PCR strip tubes and 3 μl of the appropriate mouse sample, positive control, or negative control were added. Strip tubes were spun down, then run on the appropriate PCR program as recommended by The Jackson Laboratory on a BioRad T100 Thermocycler. PCR products were run on a 2% agarose gel with 5 μl ethidium bromide per 100 ml of agarose/TAE mix. Gels were imaged using an AlphaImager Mini. Each mouse was genotyped with the Gli1-Cre, Cre, Rosa26-YFP, Myh11-Cre, and Klf4 flox (as appropriate) primer sets.

#### *In vivo* Mouse Procedures

Prior to experimentation, all mice (both on atherogenic and standard diets) were injected with tamoxifen to activate Cre recombinase. Stock tamoxifen solution was prepared using 400 mg tamoxifen (Sigma T5648-5G), 2 ml 100% ethanol, and 38 ml corn oil. The solution was incubated in a 60°C bead bath with intermittent vortexing for 3-5 hours, until the tamoxifen was fully dissolved. Tamoxifen solution was then passed through a 0.2 μm syringe filter and stored at -20°C. To activate Cre recombinase, AdvSca1-SM lineage mice between the ages of 6 and 8 weeks received 12 consecutive days of intraperitoneal injections, a dosing regimen validated by our lab’s previous research; SMC lineage mice received 7 consecutive days of injections. Each injection was 150 μl delivered through a 25G syringe (1.5 mg/day), and the side of injection was alternated daily to minimize irritation and trauma. Mice were weight stable throughout the injections (Extended Data Figure 1).

Following tamoxifen injections, mice were weighed to obtain baseline metrics. Mice randomized to the non-atherogenic arm of the study were then placed in a new group housing cage and maintained on standard chow Envigo 2920x (16% fat). Mice randomized to atherogenic treatment received a retroorbital injection of AAV bearing a mutant gain of function PCSK9 (Vector Biolabs, AAV8-D377Y-mPCSK9, Addgene 58376). PCSK9 is involved in clearance of low-density lipoprotein-cholesterol (LDL-C) from the bloodstream by mediating low density lipoprotein receptor (LDLR) internalization and lysosomal degradation. When coupled with a high fat diet, injection of 1 x 10^11^ gene copies (gc) per mouse of AAV-m-PCSK9 has been shown to induce hypercholesterolemia and atherosclerotic lesion formation in mice without ApoE-/- or LDLR-/- mutations^17^. The AAV stock was diluted in sterile saline such that each mouse received 1 x 10^11^ gc delivered in 200 μl of solution through a 28G syringe. The retroorbital injections were conducted under anesthesia per institutional protocol (induction at 3-5% isofluorane, and maintenance on the nose cone at 1.5-3% isofluorane). In a subset of mice, retroorbital blood draws were used to collect plasma at the time of PCSK9 injection and throughout the experiment. Following injection of the virus and/or retroorbital blood draws, animals were treated with Proparacaine HCl 0.5% ophthalmic eye drops (Bausch & Lomb 24208073006). Animals were observed for full recovery from the procedure, then placed in a new group housing cage and given atherogenic diet. A second dose of the AAV-m-PCSK9 was delivered after 2 weeks to ensure adequate viral load to induce hypercholesterolemia and lesion formation, and an additional dose was given at 16 and 24 weeks for mice randomized to longer endpoints. Mice were maintained under these conditions until their assigned endpoint was reached (8, 16, 24, or 28 weeks) and then were humanely euthanized.

#### Lectin Staining

A subset of mice was injected with fluorescently labeled lectin *in vivo* to identify functional vasculature. The B4 isolectin has high affinity for terminal α-D-galactosyl residues and binds to endothelial cells. The lectin solution (100 μl *Griffonia simplicifolia* lectin I (GSL I-B4), DyLight 594 + 100 μl sterile saline) was retroorbitally injected into mice (200 μl solution per mouse) and allowed to circulate for 5 minutes prior to sacrifice to label the vasculature.

#### Mouse Euthanasia and Tissue Harvest

Mice were euthanized and harvested according to institutional protocol either at their assigned endpoint or in the case of significant injury/illness. Specifically, mice were exposed to an overdose of isofluorane using the drop technique and left in the chamber until cessation of respiration. At that point, mice were removed from the chamber and cervical dislocation was performed as a secondary method of euthanasia (bilateral thoracotomy and exsanguination were also performed later during the harvest). Using a dissecting microscope, the abdominal cavity was opened, the diaphragm cut, and the ribcage removed. Blood was collected via cardiac puncture using a 25G syringe, then stored on ice for up to 1 hour during the harvest. Following exsanguination, mice were perfused with 10 ml of PBS/heparin (Sigma H3393, diluted to 0.08 KU/ml in PBS).

For tissues being collected for histology, mice were then perfused with 10 ml of 4% PFA in PBS. The heart, combined aortic arch/brachiocephalic artery (BCA)/carotid arteries, and the descending aorta were then microdissected out of the animal and placed in Eppendorf tubes containing 4% PFA. These samples were placed at 4°C overnight to ensure adequate fixation. For tissues being collected for flow cytometry or single cell RNA sequencing, the heart, aortic arch/BCA/carotids, and descending aorta (for flow cytometry only) were then microdissected out of the animal and placed in dishes containing 10% FBS MEM to maintain cell viability. These dishes were left on ice for the duration of the harvest.

#### Mouse Serum Analysis

For serum collection, tubes of blood were removed from the ice and left at room temperature for 30 minutes to clot, then the clots were dislodged using p200 pipette tips. For plasma collection, retroorbital blood draws were performed using microhematocrit capillary tubes (Fisherbrand 22-362566) and collected in heparinized microtainers (BD 365965). Blood samples were spun down in a tabletop centrifuge for 10 minutes at 4°C and 10,000 G. The supernatant (serum or plasma) was removed from the tube and transferred to a new Eppendorf for storage at -80°C until analysis.

Serum was analyzed in batches for total cholesterol to confirm that animals in the atherogenic arm of the study responded to the AAV-m-PCSK9 and high fat/high cholesterol diet. FujiFilm Wako Diagnostics kits for Cholesterol E (999-02601) were used according to manufacturer recommendations. Samples were diluted down 1:10 in sterile water so they fell on the standard curve. All tests were completed in triplicate.

#### Histology and Immunofluorescence

Tissues collected for histology were stored at 4% PFA at 4°C overnight to allow for tissue fixation. After 24 hours, tissues were removed from the PFA, rinsed with PBS, then transferred to 30% sucrose/PBS. The sucrose/PBS incubation was for a minimum of 24 hours at 4°C to ensure adequate cryoprotection. Tissues were then embedded in Tissue-Tek O.C.T. Compound (Sakura) and frozen per manufacturer recommendations. Tissue blocks were stored at -80°C until sectioning. 6 μm thick serial sections were collected from each block for use in histology and immunofluorescent/RNAscope microscopy studies.

Slides were stained with H&E and/or Masson’s Trichrome by the Anschutz Pathology Shared Resource Research Histology group. For fluorescent imaging, sections were rehydrated, permeabilized, blocked with 3% horse serum in PBS, and double or triple stained for combinations of YFP, Sca1, SMC markers (αSMA), macrophage markers (CD68), adipocyte markers (FABP44), erythrocyte markers (TER-119), and others. Slides were then imaged using a Keyence BZ-X710 microscope and BZ-X Viewer image acquisition software. For imaging, exposure and gain were consistent throughout an experiment, and signal to noise ratios were maximized. Using ImageJ/Fiji software, a minimum of two independent investigators counted cells and graded lesion size/severity in a blinded fashion.

#### RNAscope

RNAscope was used to detect colocalization of fibroblast markers (lumican) with YFP^+^ cells in the aortic root sections. Positive and negative control probes were employed. For detection of lumican, slides were prepared according to the RNAscope RED Assay and Immunofluorescence technical note (322350-TN/Rev A/Draft Date 06052017) and RNAscope 2.5 HD Detection Reagent RED User Manual (322360-USM Rev. Dat 11052015). Following RNAscope signal detection, slides were washed with PBS and stained with a FITC-conjugated anti-GFP antibody following standard immunofluorescent staining.

#### Sudan IV Staining

Sudan IV staining was used to assess lipid burden in whole aortas. The Sudan IV staining solution included 0.5 gm Sudan IV powder, 25 ml acetone, 17.5 ml 100% ethanol, and 7.5 ml DI water. Ingredients were mixed, left to rest for ∼1 hour, then filtered through a 0.2 μm Nalgene Rapid-Flow unit. The vascular tree (aortic arch through the common iliac artery) was prepared for staining by trimming fat from the perivascular area then cut open longitudinally to expose the luminal surface. Vessels were rinsed in 70% ethanol for 30 seconds, then placed in a 12 well plate and covered with Sudan IV solution. Samples were incubated at room temperature on a rocker for 30 minutes, then were rinsed twice with 80% ethanol to destain the non-plaque regions. Any remaining perivascular fat was removed, then vessels were pinned open for imaging of plaque burden.

#### Single Cell Digests

Tissues collected for flow cytometry were maintained in dishes containing 10% FBS MEM on ice for the duration of the mouse harvest to maintain cell viability. Following harvest, tissues were rinsed in HBSS to remove any serum from the samples, since serum can interfere with the tissue digest. Digest solution was made from 5mg/mL Elastase (Worthington), 0.2mg/mL Soybean Trypsin inhibitor (Sigma), 3.2mg/mL collagenase II dissolved in HBSS and filtered through a 0.2 μm syringe filter. Individual tissues were minced using dissection scissors in Eppendorf tubes with 500 μl digestion buffer, then incubated in a 37°C incubator for about 1 hour to create a single cell suspension. Samples were removed from the incubator at approximately 10-minute intervals and pipetted with a p1000 tip to facilitate tissue digestion.

At the end of the digest period, samples were spun down for 12 minutes at 172 G, 4°C to pellet the single cells. Supernatant was removed, and cells were resuspended in FA3 Buffer. The FA3 buffer was prepared with 1x PBS, 1mM EDTA, 25mM HEPES (pH 7.0), 1% FBS, then sterile filtered. Samples were centrifuged again with the same parameters, resuspended FA3, then filtered through 70 μm FLOWMI cell strainers.

#### Flow Cytometry

Following filtration, samples were spun down again, and pellets were resuspended in low volume FA3. 100 μl of each sample was placed in a 96 deep well U Bottom Plate, with excess samples pooled for use as staining controls (single color, FMO, isotype, and full stain). Plates were spun down, and 10 μl of FC block (diluted 1:10 in FA3) was added to each sample well and incubated at room temperature. After a 10-minute incubation, 100 μl of antibody cocktails for cell surface markers were added to the samples, then incubated at room temperature for 30 minutes, shielded from light. After 30 minutes, plates were spun down at 600 G (other parameters remained the same), supernatant was removed, and 150 μl Fixation Buffer (Invitrogen 88-8824-00) was added to each well. Plates were incubated at 4°C for 1 hour away from light. Plates were spun down again at 600 G, supernatant was removed, and samples were resuspended in 600 μl FA3 buffer for overnight storage. Plates were wrapped with Parafilm and aluminum foil to protect against evaporation and light exposure, then were stored at 4°C overnight.

The following day plates were spun down at 600 G, supernatant removed, and samples were rinsed with sterile PBS. After an additional spin at 600 G and decanting of supernatant, 100 μl of antibody cocktails for intracellular markers were added to each well. Plates were incubated at 4°C for 2 hours away from light. Plates were then spun again at 600 G, supernatant removed, and samples were incubated with Permeabilization Buffer (Invitrogen 88-8824-00) for 30 minutes. This process was repeated to allow for a second rinse. After the final spin, cells were resuspended in FA3 and transferred to 1.2 ml microdilution tubes. These samples were then kept at 4°C and shielded from light until ready to process on the Gallios or Gallios 561.

#### Single cell RNA sequencing (scRNA-Seq)

For single cell RNA sequencing, single cell digests were prepared from the aortic root, aortic arch, BCA, and carotid arteries as described above. To have sufficient cells for the capture/sequence, 3 mice per condition were pooled into a single sample. Following preparation of single cell digests, each sample was stained with DAPI and flow sorted to remove RBCs, cell debris, and dead cells. Cells were also sorted into YFP^+^ and YFP^-^ samples for each condition (Extended Data Figure 3). Samples were spun down and resuspended in 2% FBS PBS to a target concentration of 1,000 cells/μl. Samples were submitted to the Anschutz Genomics Shared Resource and a total of 5,000 cells per sample were captured and sequenced at a depth of 5,000 reads per cell using the 10x Genomics platform. Sequencing data were processed through the Cell Ranger pipeline with custom build reference genome (Ensembl GRCm39 release 104) containing eYFP ORF sequence. scRNA-Seq data was analyzed using Seurat and other R packages; code for this analysis has been deposited in the Weiser-Evans Lab GitHub (https://github.com/weiserevanslab/adubner). The Seurat object was analyzed with scDblFinder to detect likely cell doublets, and these were removed from the dataset (Supplementary Figure 4A). nFeature, nCount, and percent.mito were plotted and used to determine filtering criteria (nFeature_RNA > 500, nFeature_RNA < 6500, nCount_RNA > 500, nCount_RNA < 45000, percent.mito < 15), as shown in Extended Data Figure 4B-D. DimPlot of all samples showing distribution of cells from each origin sample, along with plots of time point and atherosclerosis vs. control treatment conditions, along with a confusion matrix, were used to confirm that batch effects were not observed and that no batch correction or integration was needed for this dataset (Extended Data Figure 4E-F). Seurat object was also passed through the Harmony algorithm and processed in parallel to the primary Seurat object to ensure our findings were not a result of batch effects (data not shown). Additional packages utilized in this analysis were dittoSeq to generate stacked bar plots, enrichR for GO and KEGG enrichment analysis, Ggpubr for statistical analysis of violin plots, and CellChat for inference and analysis of cell-cell communication. RNA velocity analysis was performed using scvelo in a Python environment. Top markers for each cluster can be found in Extended Data Table 1 (top genes per cluster in all conditions, all genotypes), Extended Data Table 2 (top genes per cluster in WT mice under all conditions), and Extended Data Table 3 (top genes per cluster in KO mice under all conditions).

### QUANTIFICATION AND STATISTICAL ANALYSIS

All experimental data were evaluated for normality through comparison of median and means, standard deviations, and through formal normality tests (Shapiro-Wilk). The appropriate statistical analysis was used, dependent on the outcome of the normality analyses. Normally distributed continuous data were analyzed using Student’s T-test, whereas skewed data were analyzed using the non-parametric Mann-Whitney U. α level was set to 0.05 and statistical tests were completed using GraphPad Prism 9; statistical analysis of violin plots was completed using the Ggpubr R package. Bar graph data was plotted as mean plus standard deviation, with all individual values displayed. Analysis of scRNA-seq data was done in conjunction with the Anschutz RNA Biosciences Initiative and Dr. Sizhao Lu.

### KEY RESOURCES TABLE

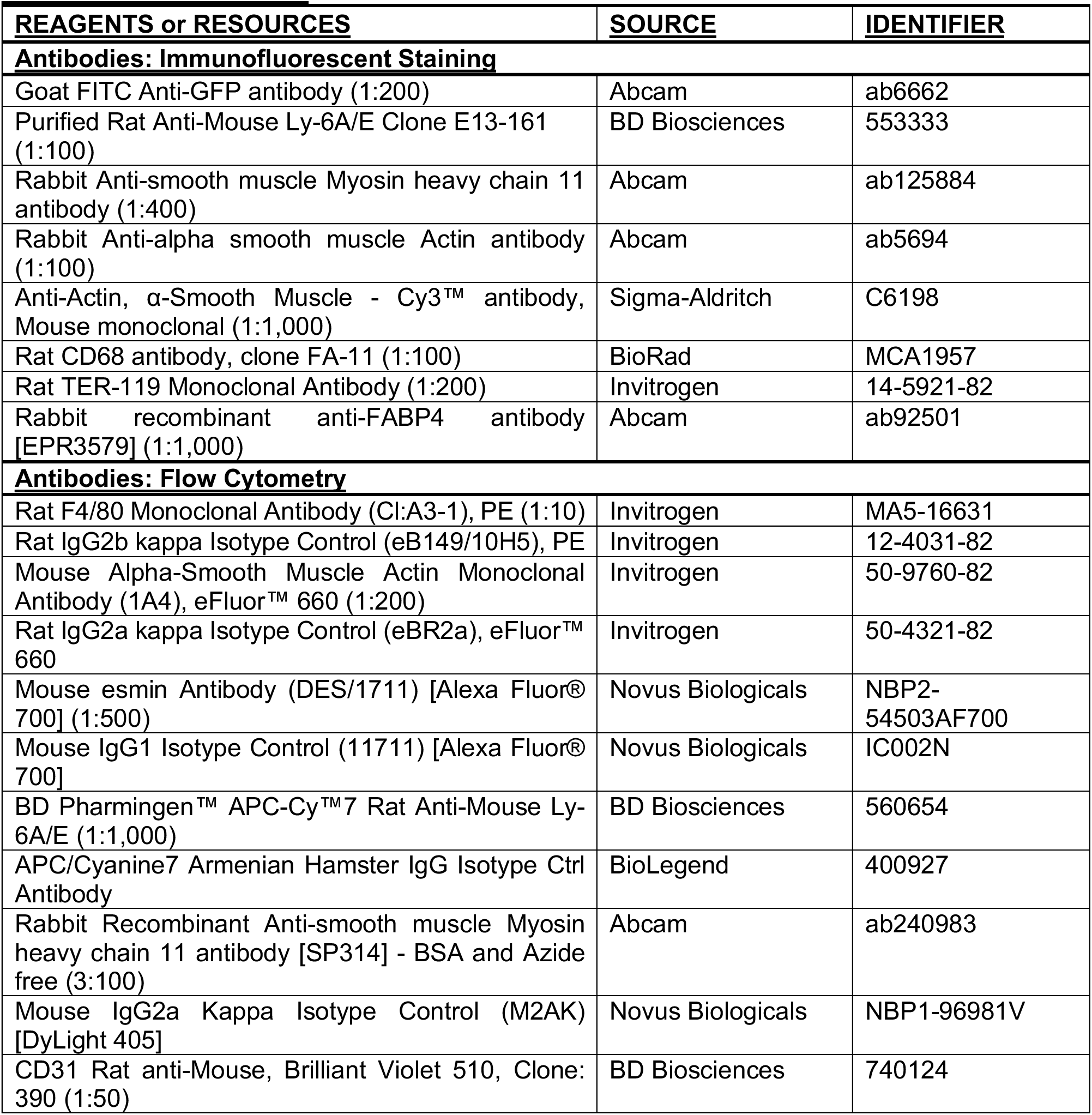
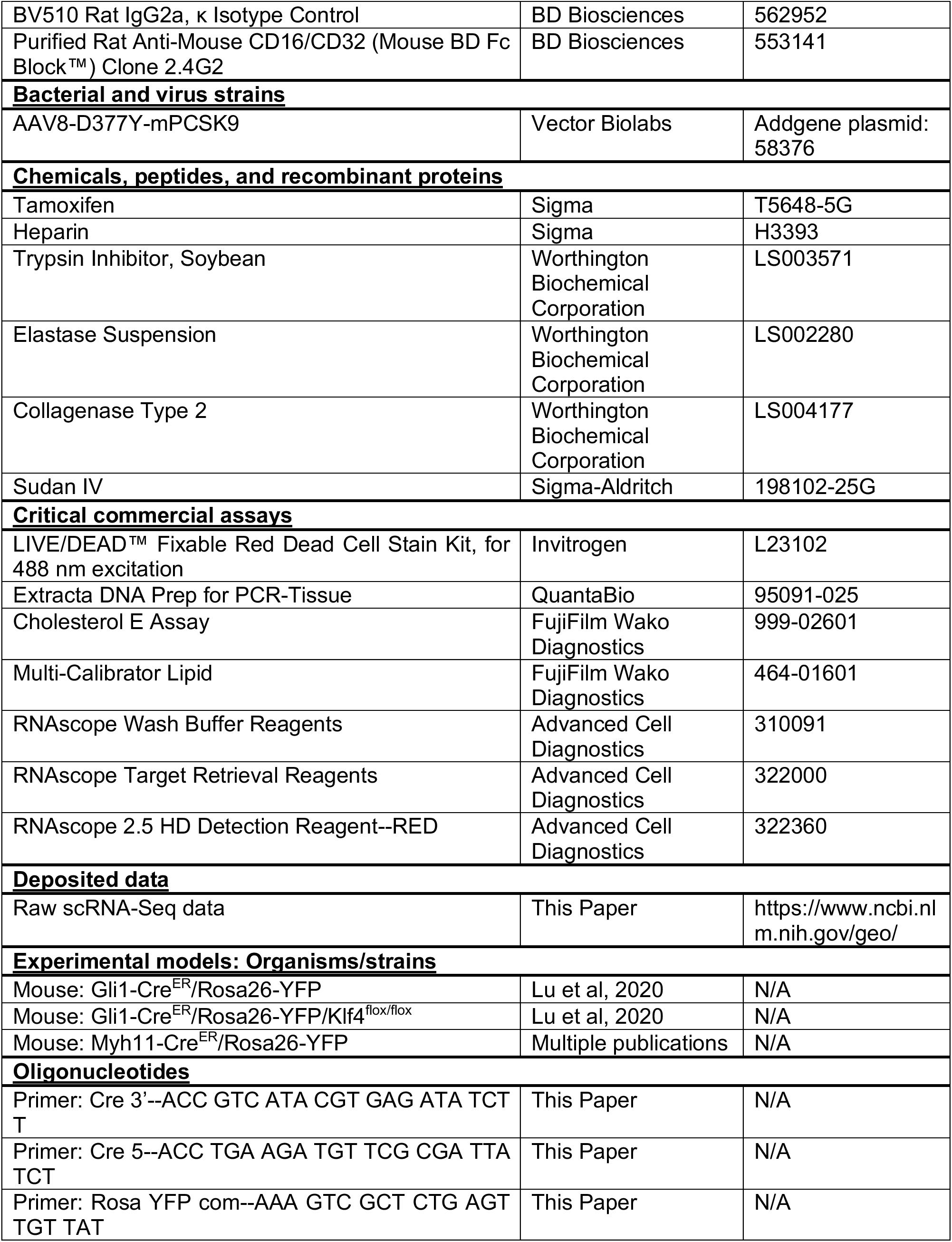
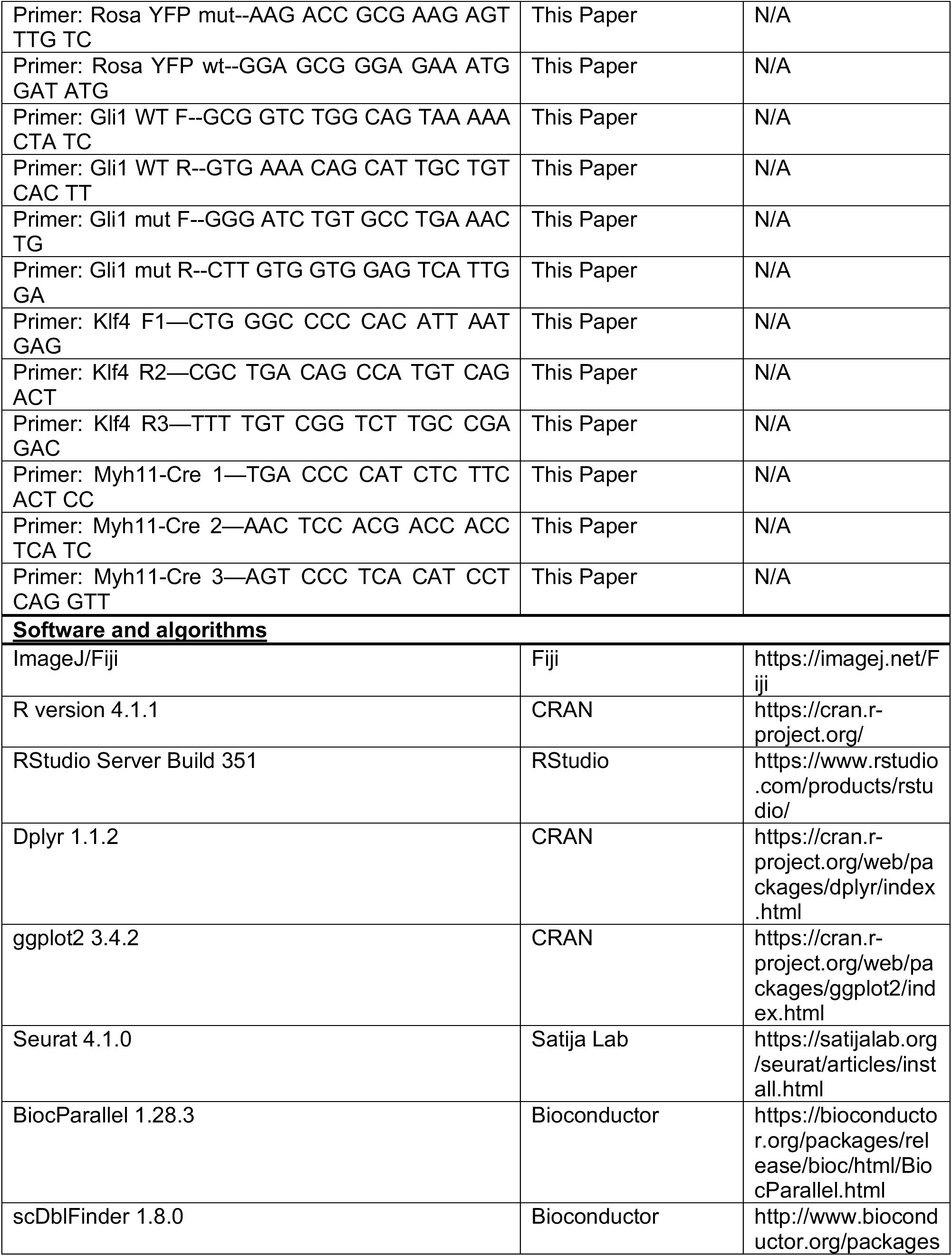
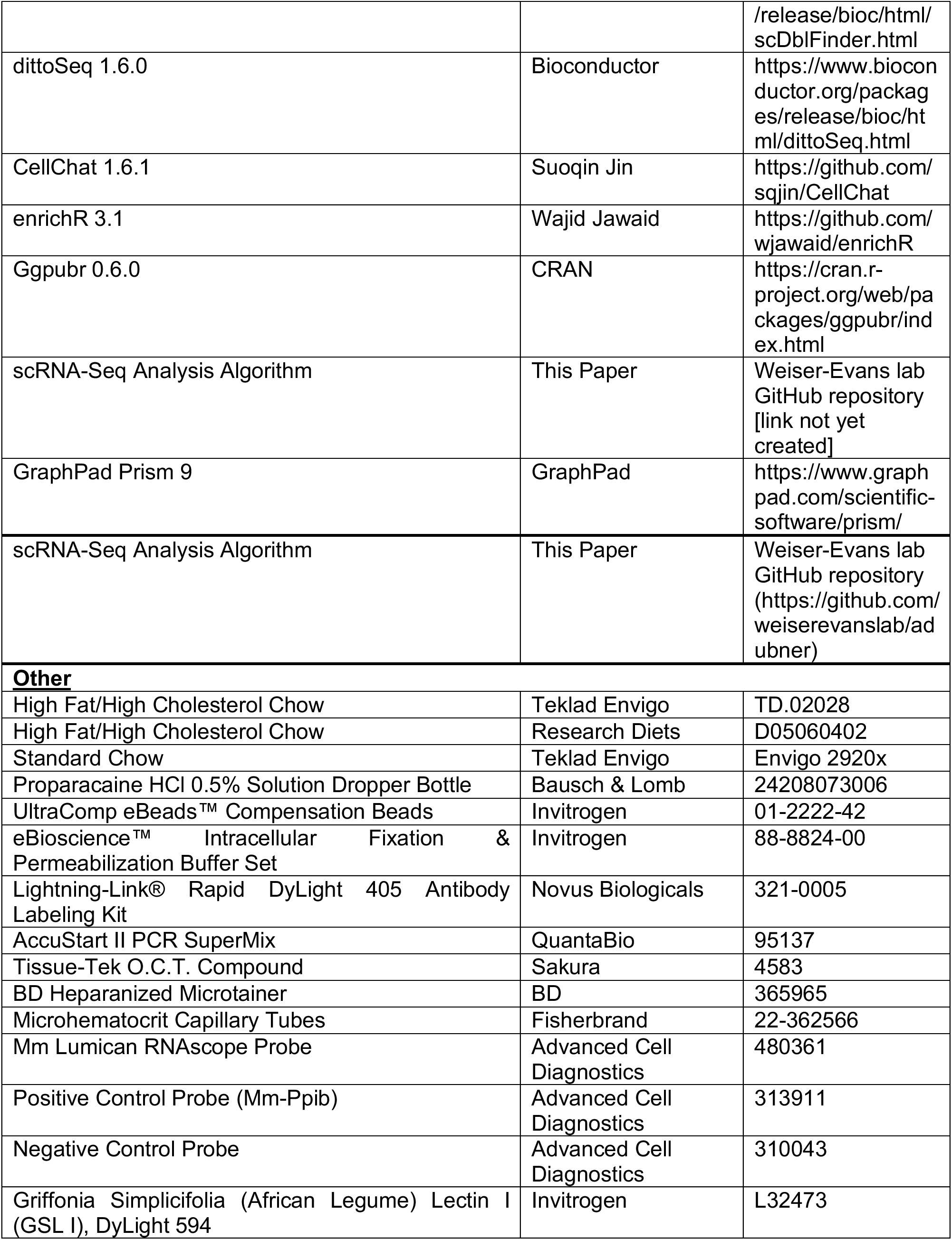

## ACKNOWLEDGEMENTS

This work was supported by grants R01 HL 121877 (Weiser-Evans, Majesky), R01 HL151331 (Weiser-Evans), R01 HL148167 (Kovacic; Weiser-Evans subcontract), and F31 HL160149-01 (Dubner) from the National Heart, Lung, and Blood Institute as well as TL1TR002533 (Dubner) from the National Center for Advancing Translational Sciences. Flow cytometry experiments were performed at the University of Colorado Cancer Center Flow Cytometry Shared Resource Core supported by Cancer Center Support Grant (no. P30CA046934). We would like to thank Dmitry Baturin for his assistance with flow sorting and Christine Childs for her assistance with the Gallios flow cytometry. We would like to thank E. Erin Smith of the University of Colorado Cancer Center Pathology Shared Resource for her assistance with histological staining. This resource is supported in part by the Cancer Center Support Grant (no. P30CA046934). The scRNA-Seq experiments were performed in conjunction with the University of Colorado Cancer Center Genomics and Microarray Shared Resource supported by Cancer Center Support Grant (no. P30CA046934), and we would like to thank Okyong Cho for all her help. We greatly appreciate the assistance of the Anschutz RNA Bioscience initiative, and particularly Caitlin Winkler, in the analysis of the scRNA-Seq datasets. We also would like to thank Andrew Glugla for his contribution of equipment for the computational analyses.

