## Supplementary material for "Smooth muscle-derived adventitial progenitor cells promote key cell type transitions controlling plaque stability in atherosclerosis in a Klf4-dependent manner": Supplemetal Data

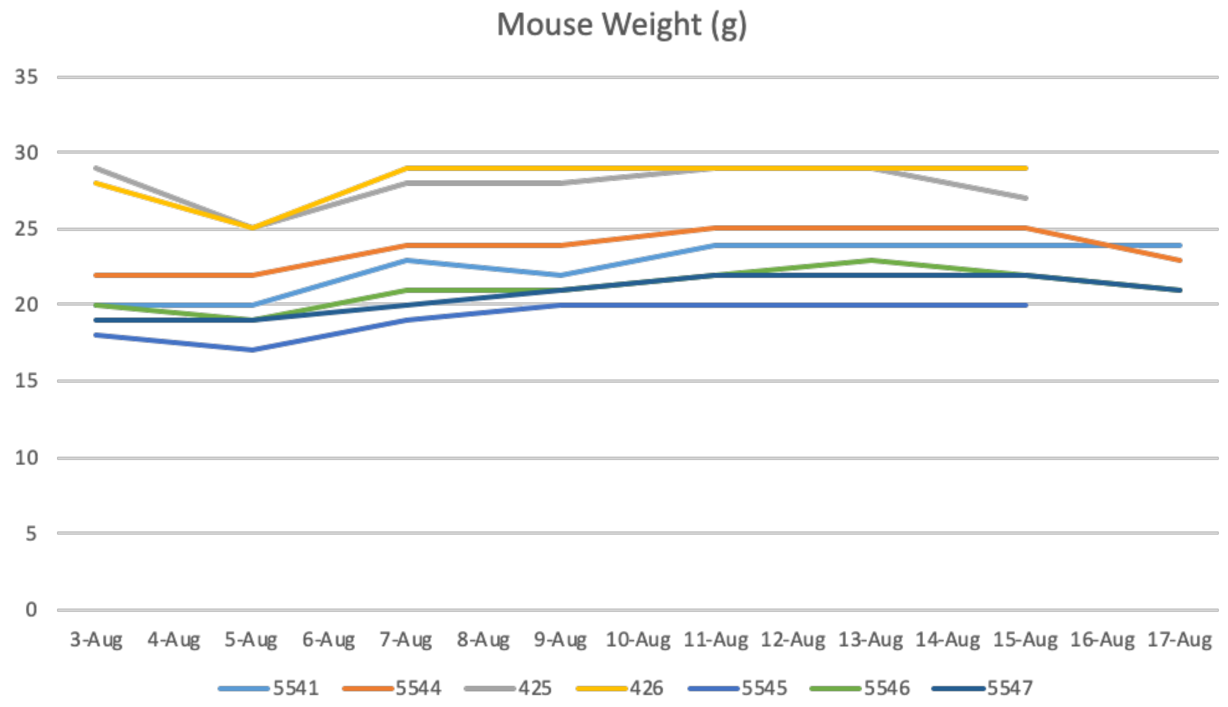


**Extended Data Figure 1. Mice are weight stable with 12 days of IP tamoxifen injections.** To induce YFP reporter knock in, mice were subjected to 12 days of IP tamoxifen dissolved in corn oil (1.5 mg). During this time, mice experienced a slight drop in weight after the first injections, but then recovered lost weight and remained weight stable for the duration of the injections.

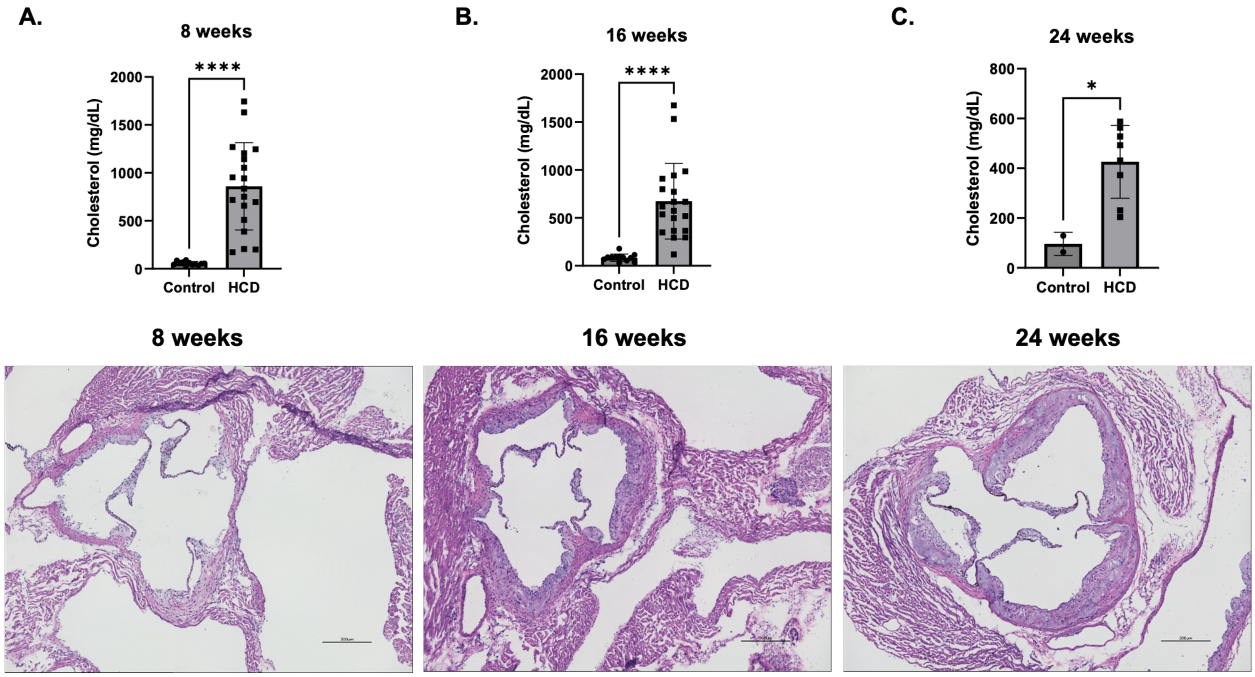


**Extended Data Figure 2. PCSK9 induces hypercholesterolemia and plaque formation. A-C.** Student's t tests (8, 24 weeks) or Mann Whitney U test (16 weeks) show that after 8, 16, and 24 weeks, mice on high fat/high cholesterol diet exhibited significantly higher serum cholesterol compared to control mice on standard chow ($p < 0.0001$). Representative H&Es show early plaque formation (8 weeks), mid-stage plaque formation (16 weeks), and late-stage plaque formation (24 weeks) in the aortic root.

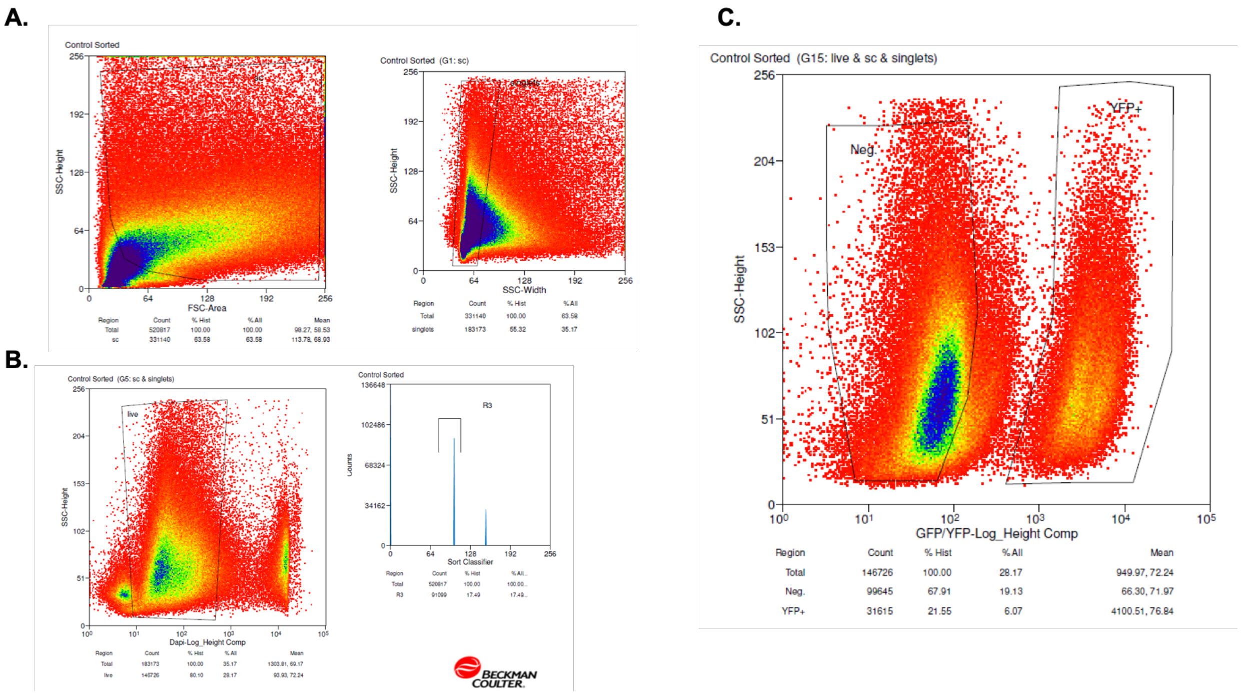


**Extended Data Figure 3. FACS sorting for YFP expression. A.** Prior to single cell RNA sequencing, single cell digests were FACS sorted. Representative image of gating strategy using side scatter and forward scatter to exclude red blood cells and cell debris. **B.** Representative image of gating strategy using DAPI staining to ensure a live cell population. **C.** Representative image of gating strategy for $YFP^+$ and $YFP^-$ cell populations, which then were captured and sequenced separately.

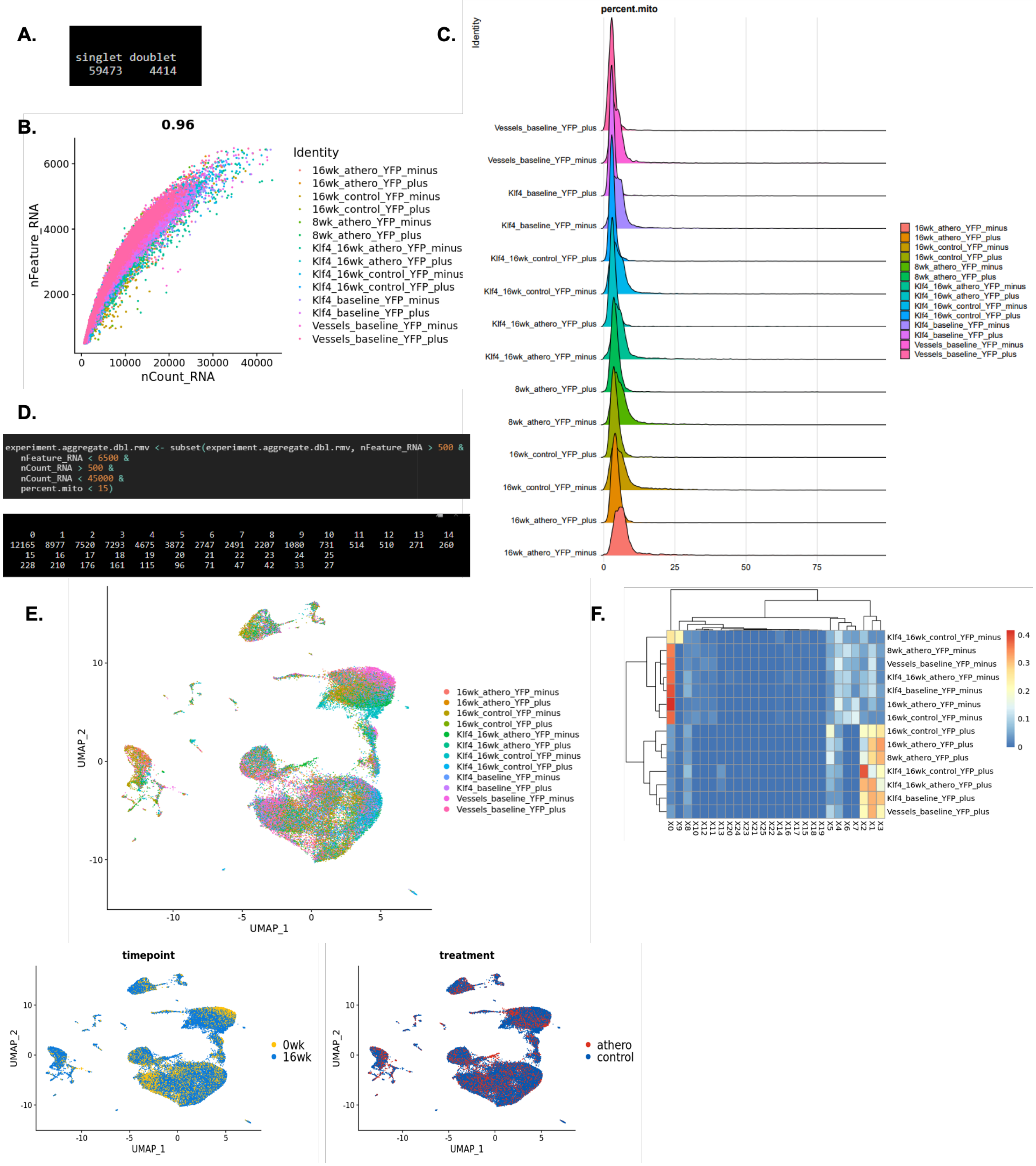


**Extended Data Figure 4. scRNA-Seq quality control and clustering. A.** The scDblFinder package in Seurat was used to analyze data and detect likely doublets. 4,414 doublets were detected and removed from the dataset. **B.** Feature Scatter plot of nFeature vs. nCount for all cells (n = 59,473). **C.** Plot of percent mitochondrial genes on average in each sample. **D.** Data was subsetted for nFeature < 6500, nCount > 500, nCount < 45000, and percent.mito <15. After continuing through the Seurat pipeline (data normalization, identification of variable genes, and scaling data), clustering at a resolution of 0.5 resulted in a total of 26 clusters. **E.** DimPlot of all samples showing distribution of cells from each origin sample, along with plots of time point and atherosclerosis vs. control treatment conditions. **F.** Confusion matrix was used to confirm the lack of batch effects in our experiments, therefore no batch correction or integration was used.

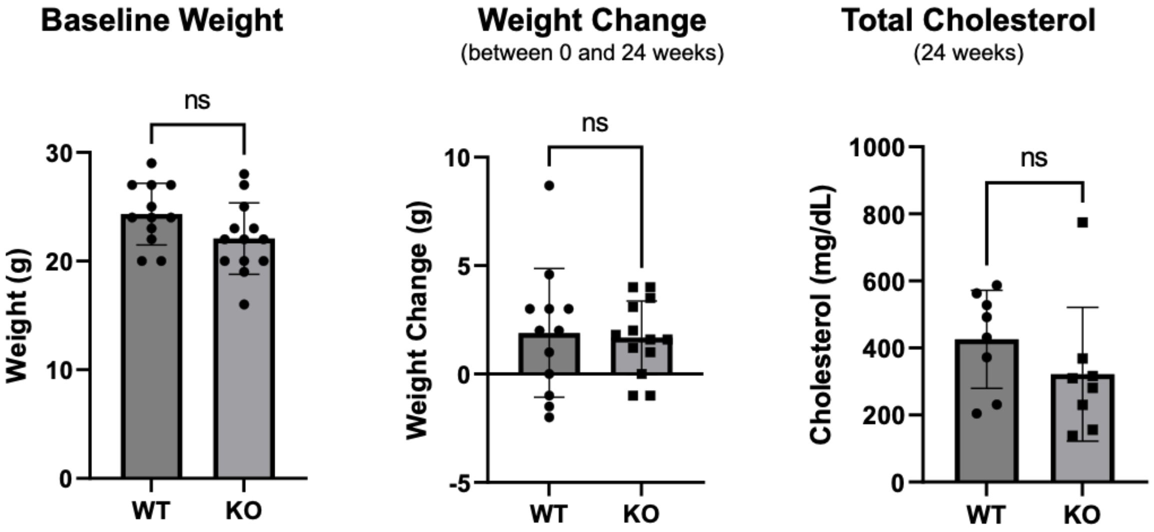


**Extended Data Figure 5. No differences in weight or total serum cholesterol between WT and Klf4 KO mice.** Student's t-test showed no difference between WT and Klf4 KO weight at baseline ($p = 0.0741$) or weight change during the 24-week experiments ($p = 0.8174$). No differences were observed with total cholesterol levels between WT and KO mice.

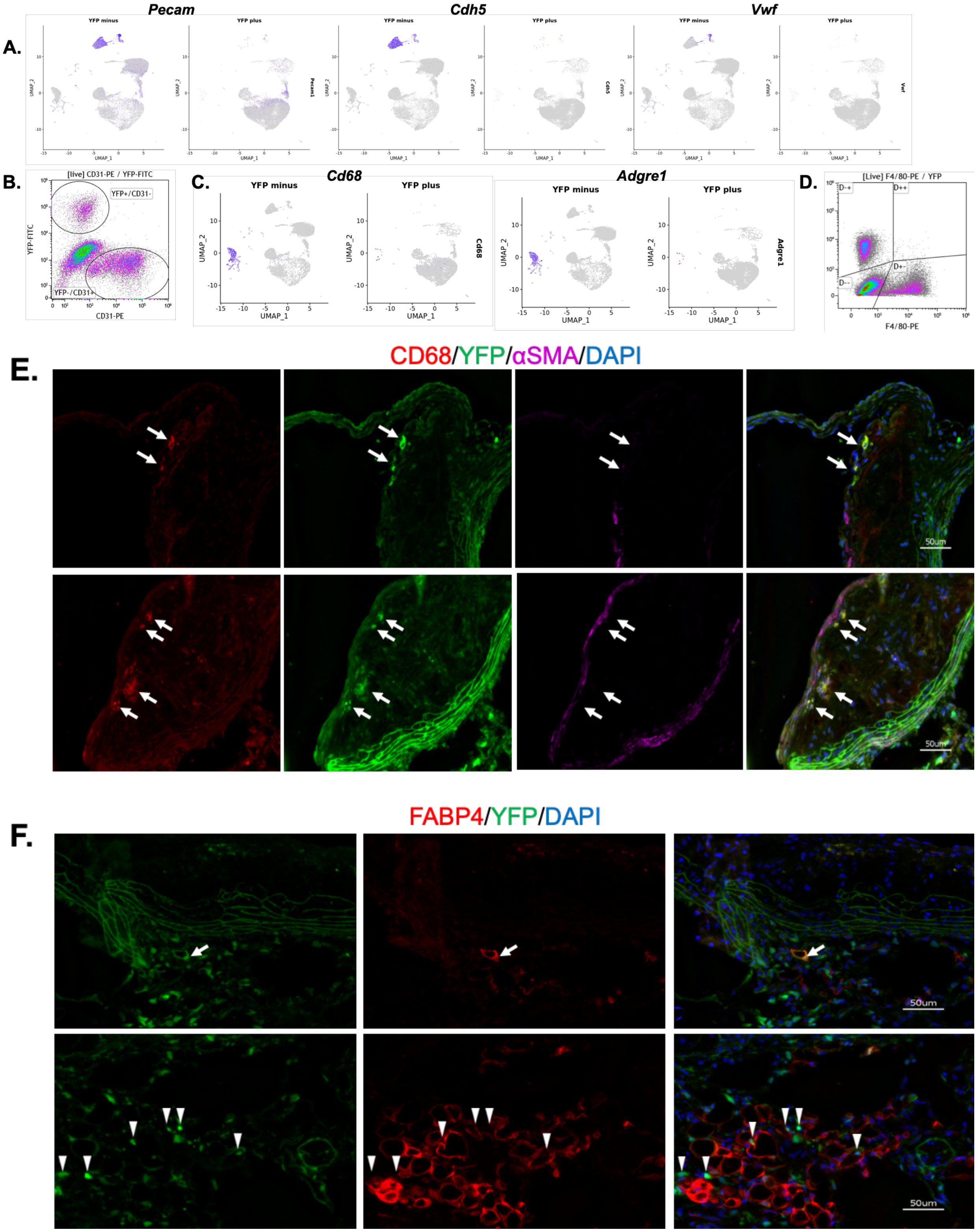


**Extended Data Figure 6. Rare differentiation trajectories for AdvSca1-SM cells.** Our experiments identified some of the rarer differentiation patterns for these AdvSca1-SM progenitor cells. **A.** Feature plots demonstrating a small number of YFP$^{+}$ cells in the endothelial cell clusters, where they are positive for endothelial cell markers (*Pecam1, Cdh5, Vwf*). **B.** Representative image of flow cytometry data after 16 weeks of atherogenic diet plotting YFP against endothelial marker CD31, showing a small number of double positive cells. **C.** Feature plots demonstrating a small number of $YFP^+$ cells in the macrophage cell clusters, where they are positive for macrophage markers (*Cd68, Adgre1*). **D.** Representative image of flow cytometry data after 16 weeks of atherogenic diet plotting YFP against macrophage marker F4/80, showing a small number of double positive cells. **E.** Representative immunofluorescent staining of aortic root plaques after 24 weeks of atherogenic treatment. CD68 (red), YFP (green), aSMA (magenta), DAPI (blue). Arrows indicate $YFP^+/CD68^+$ cells that are $\alpha SMA^-$. **F.** Representative immunofluorescent staining of aortic root plaques after 24 weeks of atherogenic treatment. Arrows indicate $YFP^+/FABP4^+$ cells. FABP4 (red), YFP (green), aSMA (magenta), DAPI (blue). Arrowheads indicate $YFP^+$ structures in proximity to $Fabp4^+$ adipocytes.

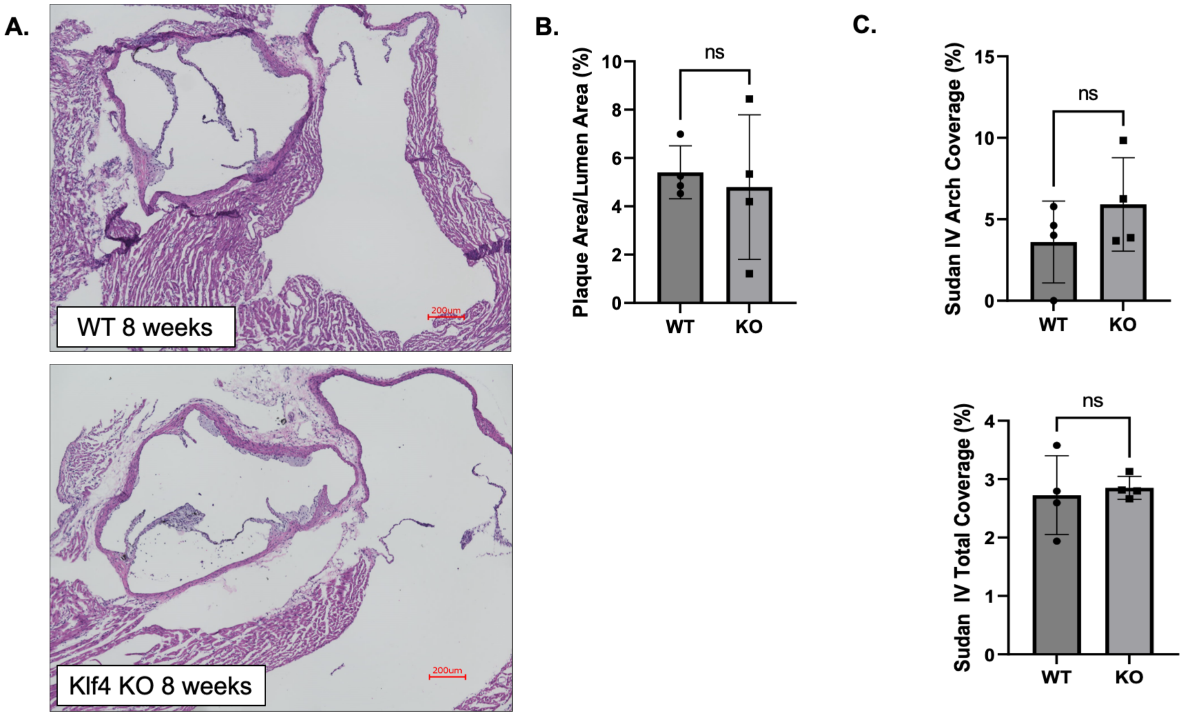


**Extended Data Figure 7. No difference between WT and KO animals in early-stage plaques.** **A.** Early-stage aortic root plaques (8 weeks atherogenic treatment) were stained with H&E to view plaque formation. **B.** Student's t-test showed no difference between plaques from WT (n = 4) and Klf4 KO (n = 4) mice in overall plaque area (data not shown) or percent coverage of the aortic root (p = 0.7517). **C.** Student's t-test showed no difference between WT (n = 4) and Klf4 KO (n = 4) in Sudan IV staining within the aortic arch (p = 0.272) or the entire vascular tree (p = 0.7322).

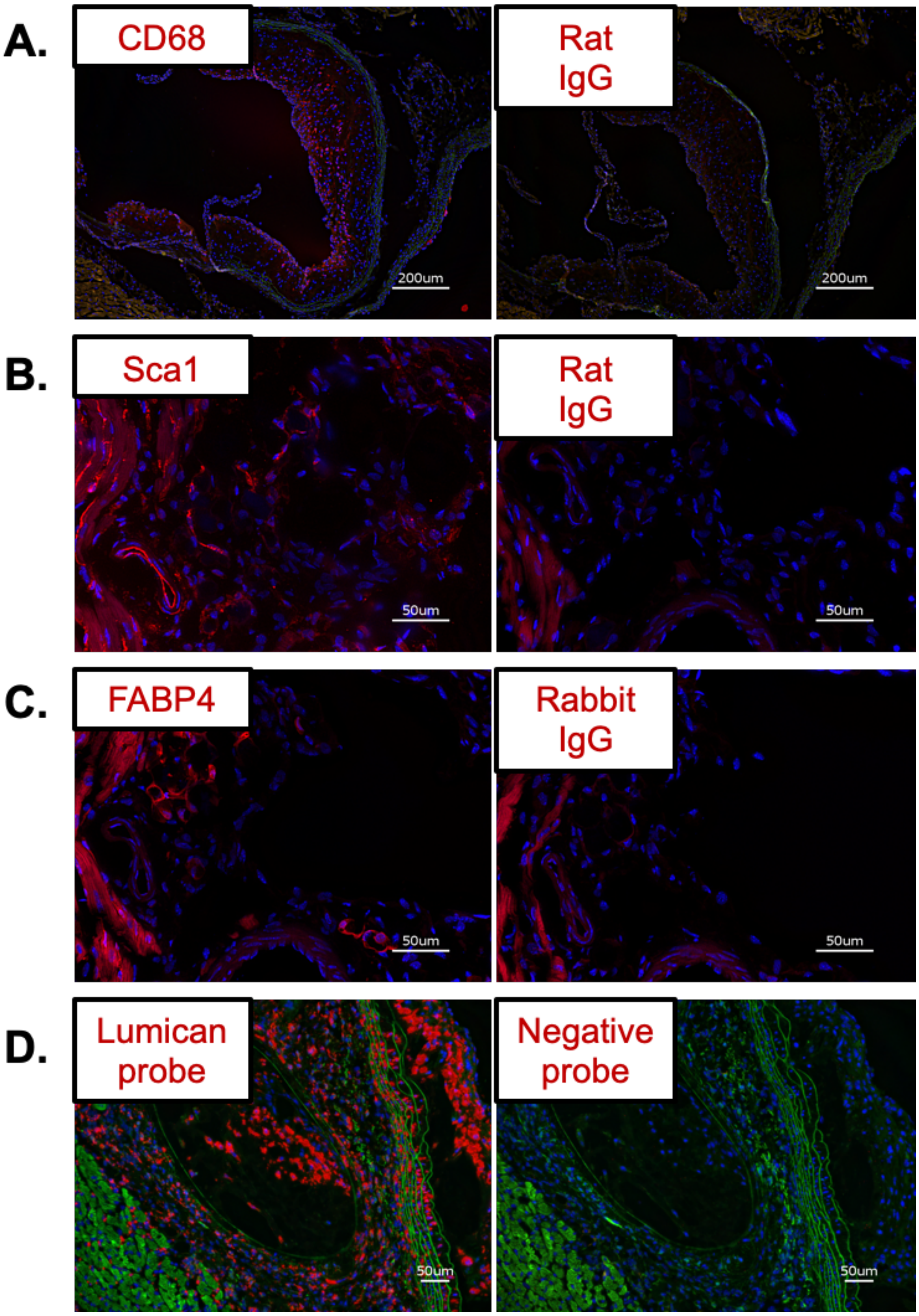


**Extended Data Figure 8: Negative staining controls**. Representative images of CD68 (**A.**), Sca1 (**B.**), FABP4 (**C.**), and Lumican ISH (**D.**) staining along with their appropriate negative controls. All negative controls were used at the same concentrations as the antibodies/probes, and exposure times were kept constant when imaging.
